# Reconfigurable asymmetric protein assemblies through implicit negative design

**DOI:** 10.1101/2021.08.15.456388

**Authors:** Danny D. Sahtoe, Florian Praetorius, Alexis Courbet, Yang Hsia, Basile I.M. Wicky, Natasha I. Edman, Lauren M. Miller, Bart J. R. Timmermans, Hana M. Morris, Alex Kang, Asim K. Bera, David Baker

## Abstract

Asymmetric multi-protein complexes that undergo subunit exchange play central roles in biology, but present a challenge for protein design. The individual components must contain interfaces enabling reversible addition to and dissociation from the complex, but be stable and well behaved in isolation. Here we employ a set of implicit negative design principles to generate beta sheet mediated heterodimers which enable the generation of a wide variety of structurally well defined asymmetric assemblies. Crystal structures of the heterodimers are very close to the design models, and unlike previously designed orthogonal heterodimer sets, the subunits are stable, folded and monomeric in isolation and rapidly assemble upon mixing. Rigid fusion of individual heterodimer halves to repeat proteins yields central assembly hubs that can bind two or three different proteins across different interfaces. We use these connectors to assemble linearly arranged hetero-oligomers with up to 6 unique components, branched hetero-oligomers, closed C4-symmetric two-component rings, and hetero-oligomers assembled on a cyclic homo-oligomeric central hub, and demonstrate such complexes can readily reconfigure through subunit exchange. Our approach provides a general route to designing asymmetric reconfigurable protein systems.

## Main Text

Dynamic reconfigurable multi-protein complexes play key roles in DNA replication, transcription, protein degradation and other central biological processes (*1*). Because the subunits of these complexes are well behaved proteins on their own, the assemblies can reconfigure throughout complex processes by successive addition or removal of one or more components. Such modulation is essential to their function: for example, orderly progression from initiation to elongation brought about by subunit loss and addition underlies the molecular mechanisms of protein complexes that drive DNA replication and transcription (*2, 3*).

The ability to de novo design such multicomponent reconfigurable protein assemblies would enable the creation of more sophisticated new functions than currently possible. There has been considerable progress in designing proteins which assemble into cyclic oligomeric and higher order symmetric nanostructures such as icosahedral nanocages with as many as 120 subunits, and 2D-layers with many thousands of regularly arrayed components (*4–8*). Essential to the ability to accurately design such systems are the symmetry and cooperativity of assembly, which result in the strong favoring of just one of the very large number of possible states. Once formed, these assemblies are therefore typically quite static and exchange subunits only on very long time scales, which is advantageous for applications such as nanoparticle vaccine design and multivalent receptor engagement ( *9*).

The design of reconfigurable asymmetric assemblies is a more difficult challenge, as there is no symmetry “bonus” favoring the target structure (as is attained for example in the closing of an icosahedral cage), and because the individual subunits must be stable and soluble proteins in isolation in order to reversibly associate or dissociate. Reconfigurable asymmetric protein assemblies could in principle be constructed using a modular set of protein-protein interaction pairs (heterodimers), provided first, that the interaction pairs are specific, second, that individual components are stable both in isolation and in complex so they can be added and removed, and third, that they can be rigidly fused to other components without changing the dimerization properties. Rigid fusion, as opposed to fusion by flexible linkers, is important to program the assembly of structurally well defined complexes, as most higher order natural protein complexes have, despite their reconfigurability, distinct overall shapes that are critical for their function.

While there has been considerable progress in the design of orthogonal sets of interacting proteins that have one of these properties, designed proteins having all of these properties are still lacking. Large sets of orthogonally interacting helical hairpin-based heterodimers have been designed by incorporating designed hydrogen bonds across extensive interaction interfaces ( *10, 11*). However, the individually expressed components in these systems form homodimers or other higher order homomeric aggregates that disassemble on very long time scales or not at all (*10, 12*), making them unsuitable for use in constructing reconfigurable higher order assemblies. Designed heterodimeric coiled-coils assemble from peptides that, in isolation, are soluble and monomeric, but the monomers are unfolded prior to binding their partners (*13, 14*), complicating their use in structurally defined rigid fusions.

We set out to design sets of interacting protein pairs with properties required for subsequent programming of reconfigurable protein assemblies (Fig. 1A). The first challenge to overcome is the systematic design of proteins with interaction surfaces that drive association with cognate partners, but not self association. This is not straightforward, as hydrophobic interactions provide a driving force for protein assembly, but these same hydrophobic residues can then mediate undesired self-self interactions. Previously designed heterodimeric helical bundles featured, in addition to hydrophobic interactions, explicit hydrogen bond networks that contribute to binding specificity and make the interface more polar. However, the individual protomers, either helical hairpins or individual helices, lack a hydrophobic core and are thus flexible and unstable in isolation, allowing a wide range of potential off-target homo-oligomers to form (Fig. 1B). Explicit negative design methods have been developed which allow the favoring of one state over another by considering the effect of amino acid substitutions on the free energies of both states ( *15, 16*). However, such methods cannot be readily applied to the general problem of disfavoring self association, as there are in general a very large number of possible self associated states which cannot be systematically enumerated.

**Fig. 1.**
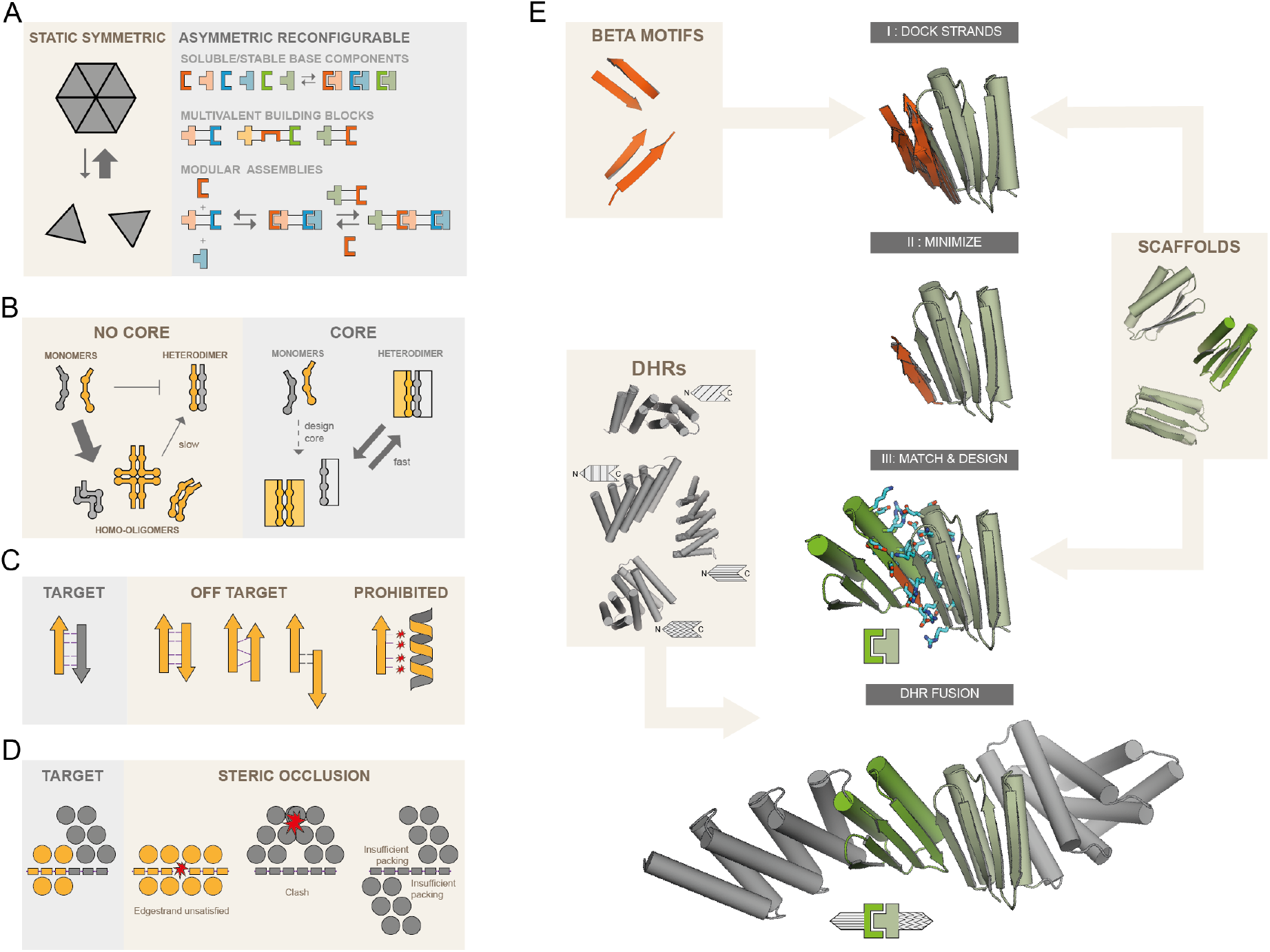
Strategies for the design of asymmetric hetero-oligomeric complexes. **(A)** Many design efforts have focused on cooperatively assembling symmetric complexes (left) with little subunit exchange. Here instead we sought to create asymmetric hetero-oligomers from stable heterodimeric building blocks, that can modularly exchange subunits (right). **(B,C,D)** Schematic illustration of properties that can contribute to prevent self-association. **(B)** Protomers that have a substantial hydrophobic core (right rectangles) are less likely to form stable homo-oligomers than protomers of previously designed heterodimers lacking hydrophobic monomer cores. **(C)** In beta-sheet extended interfaces, most homodimer states that bury non h-bonding polar edge strand atoms are energetically inaccessible. Potential homodimers are more likely to form via beta sheet extension. These are restricted to only 2 orientations (parallel and antiparallel) and a limited number of offset registers. Arrows and ribbons represent strands and helices, respectively; thin lines indicate hydrogen bonds, red stars indicate unsatisfied polar groups. **(D)** “Cross sectional” schematic view (helices as circles, beta strands as rectangles, star indicates steric clash) By modeling the limited number of beta sheet homodimers across the beta edge strand, structural elements may be designed that specifically block homodimer formation but still allow heterodimer formation. **(E)** Design workflow: Beta sheet motifs are docked to the edge strands of a library of hydrophobic core containing fold-it scaffolds. Minimized docked strands are incorporated into scaffolds by matching the strands to the scaffold library, yielding docked protein-protein complexes, followed by interface sequence design. Resulting docks are fused rigidly on their terminal helices to a library of DHRs.

We instead sought to use implicit negative design (*17*) by introducing three properties that collectively make self associated states unlikely to have low free energy: First, in contrast to the flexible coiled coil and helical hairpins in previous designs, we aimed for well folded individual protomers stabilized by substantial hydrophobic cores; this property limits the formation of slowly-exchanging homo-oligomers (Fig. 1B). Second, we constructed interfaces in which each protomer has a mixed alpha-beta topology and contributes one exposed beta strand to the interface, giving rise to a continuous beta sheet across the heterodimer interface ( *18–20*) (Fig. 1C). The exposed polar backbone atoms of this “edge strand” limit undesired self-association to arrangements that pair the beta edge strands; most other homomeric arrangements result in energetically unfavorable burial of the polar backbone atoms on the beta edge strand and hence are unlikely to form (Fig. 1C). Third, we incorporated structural elements likely to clash in undesired homomeric states (steric occlusion). The restrictions in possible undesired states resulting from strategies 1 and 2 make it possible to explicitly model the limited number of homo-oligomeric states, and hence to explicitly design in additional elements likely to sterically occlude such states (Fig. 1D).

To implement these properties in actual proteins, we chose to start with a set of mixed alpha/beta scaffolds that were designed by FoldIt players (*21*). The selected designs contain sizable hydrophobic cores, exposed edge strands required for beta sheet extension (*18*) and one terminal helix as needed for rigid helical fusion (Fig. 1E) (*22*). Using blueprint-based backbone building (*23, 24*) we designed additional helices at the other terminus for a subset of the scaffolds to enable rigid fusion at both the N and C termini (Fig. S1). Heterodimers with beta sheets extending across the interface were generated by superimposing one of the two strands from each of a series of paired beta strand templates on an edge beta strand of each scaffold (Fig. 1E, top), and then optimizing the rigid body orientation and the internal geometry of the partner beta strand to maximize hydrogen bonding interactions across the interface (Fig. 1E, second row). This generates a series of disembodied beta strands forming an extended beta sheet for each scaffold; for each of these, an edge beta strand from a second scaffold was superimposed on the disembodied beta strand to form an extended sheet-on-sheet interface (Fig. 1E, third row). The interface sidechain-sidechain interactions in the resulting protein-protein docks were optimized using Rosetta combinatorial sequence design (*25*). To limit excessive hydrophobic interactions, we either generated explicit hydrogen bond networks across the heterodimer interface ( *11*), or used compositional constraints to encourage the use of polar residues while penalizing buried unsatisfied polar groups (*26*). This resulted in interfaces that, outside of the polar hydrogen bonding of the beta strands, contained both hydrophobic interactions and polar networks. To further disfavor unwanted homodimeric interactions (Fig. 1D, right panel), and to facilitate incorporation of the heterodimeric building blocks into higher order assemblies, we rigidly fused designed helical repeat proteins (DHRs) to terminal helices (*22, 27*). Designed heterodimers were selected for experimental characterization based on binding energy, the number of buried unsatisfied polar groups, buried surface area and shape complementarity (see methods).

We co-expressed the selected heterodimers in *E. coli* using a bicistronic expression system encoding one of the two protomers with a C-terminal polyhistidine tag and the other either untagged or GFP-tagged at the N-terminus. Complex formation was initially assessed using nickel affinity chromatography; designs for which both protomers were present in SDS-PAGE after nickel pulldown were subsequently subjected to size exclusion chromatography (SEC) and liquid chromatography - mass spectrometry (LC/MS). Of the 238 tested designs, 71 passed the bicistronic screen and were selected for individual expression of protomers. Of these, 32 formed heterodimers from individually purified monomers as confirmed by SEC, native MS, or both (Fig. 2A, Fig. S2). In SEC titration experiments, some protomers were monomeric at all injection concentrations, while others self-associated at higher concentrations (Fig. S3). Both LHD101 protomers and their fusions were monomeric even at injection concentrations above 100 µM (Fig. S3). LHD275A, LHD278A, LHD317A, and a redesigned version of LHD29 with a more polar interface (LHD274) were also predominantly monomeric (Fig. S3; Fig. S4). Designs for which isolated protomers were poorly expressed, polydisperse in SEC or did not yield stable, soluble and functional rigid DHR fusions were discarded together with designs that were very similar to other designs, but otherwise behaved well. After this stringent selection, we were left with a set of 11 heterodimers spanning three main structural classes (Fig. 2A, Fig. S2A). In class one, the central extended beta sheet is buttressed on opposite sides by helices that contribute additional interface interactions (LHDs 29 and 202 in Fig. 2A), in class two the helices that provide additional interactions are on the same side of the extended central sheet (LHDs 101 and 206 in Fig. 2A), and in the third class, both sides of the central beta sheet extension are flanked by helices (LHDs 275 and 317 in Fig. 2A).

**Fig 2.**
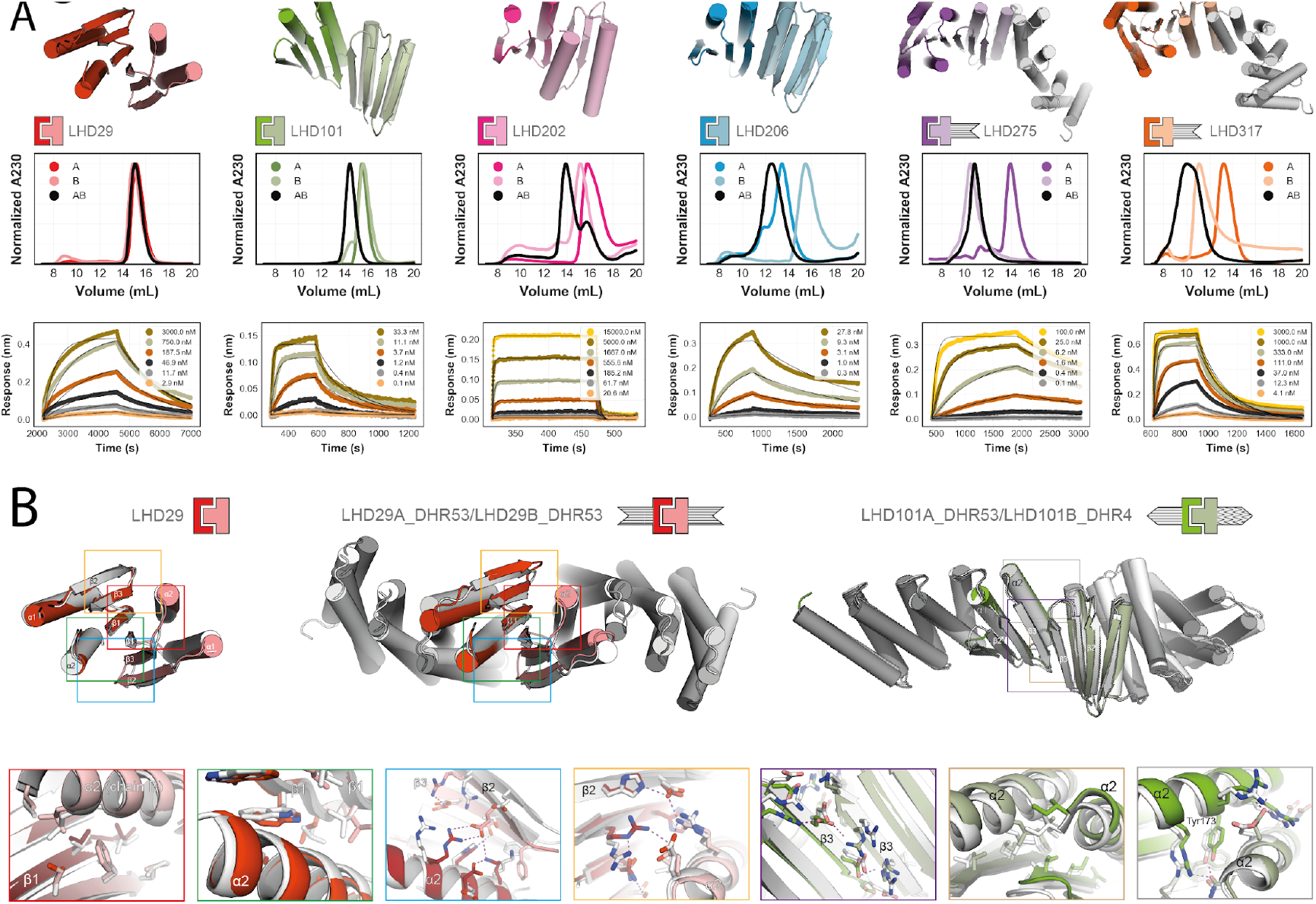
Experimental characterization. **(A)** Top row, design models of six different heterodimers. Middle row, normalized SEC traces of individual protomers (A, B) and complexes (AB). Bottom row, kinetic binding traces with global kinetic fits of in vitro biolayer interferometry binding assays. **(B):** Crystal structures (in colors) of the designs LHD29, LHD29A53/B53 and LHD101A53/B4 overlayed on design models (light gray). Colored rectangles in the full models (top row) match the corresponding detailed views (bottom row).

We monitored the kinetics of heterodimer formation and dissociation through biolayer interferometry (BLI) (Fig. 2A, Fig. S2A,C and table S1) by immobilizing individual biotinylated protomers onto streptavidin coated sensors and adding the designed binding partner. Unlike previously designed heterodimers, binding reactions equilibrated rapidly. Differences in off rates indicate that the heterodimers span a range of affinities (Fig. S2D and table S1). Association rates were quite fast and ranged from 10^6^ M^-1^ s^-1^ for the fastest heterodimer to 10^2^ M^-1^ s^-1^ for the slowest heterodimer LHD29; even LHD29 equilibrated an order of magnitude faster than the fastest associating designed helical hairpin heterodimer (Fig. 2A, Fig. S5A, Table S2). For LHD101 and LHD206 we confirmed BLI measurements in a split luciferase-based binding assay performed in E.coli lysates. The Kd’s agreed well with those from BLI, showing that heterodimer association is not affected by high concentrations of non-cognate proteins (Fig. S5D,E and Table S3).

We determined the crystal structures of two class one designs, LHD29 (2.2 Å) and LHD29A53/B53 (2.6 Å) in which both protomers are fused to DHR53 (Fig 2B and table S4). In the central extended beta sheet, the LHD29 design closely matches the crystal structure (Fig. 2B, red and green box). Aside from backbone beta sheet hydrogens bonds, this part of the interface is supported by primarily hydrophobic packing interactions between the side chains of each interface beta edge strand. The two flanking helices on opposite sides of the central beta sheet (Figure 2B blue and orange box) contribute predominantly polar contacts to the interface, and are also very similar in the crystal structure and design model. Apart from crystal contact induced subtle backbone rearrangements in strand 2 of LHD29B, that promote the formation of a polar interaction network (Figure 2B blue box), most interface sidechain-sidechain interactions agree well with the design model. Similar to the unfused LHD29, the interface of LHD29A53/B53 closely resembles the designed model; at the fusion junction and repeat protein regions, deviations are slightly larger.

We also determined the structure of a class two design, LHD101A53/B4 (2.2 Å), in which protomer A is fused to DHR53 and B to DHR4 (Fig. 2B and table S4). The crystal structure is again very close to the design model at both the interface and fusion junction, as well as the repeat protein region. In class two designs, the interface beta strand pair is reinforced by flanking helices that, unlike class one designs, are in direct contact with both each other and the interface beta sheet. The solvent exposed side of the beta interface consists primarily of electrostatic interactions (Fig. 2B, purple box). The buried side of the beta interface consists of exclusively hydrophobic side chains. Together with apolar side chains on the flanking helices of both protomers, these residues form a closely packed core interface (Fig. 2B, brown box) that is further stabilized by solvent exposed polar interactions between the flanking helices. Notably, the designed semi-buried polar interaction network centered on Tyr173 is maintained in the crystal structure (Fig. 2B, gray box).

As described above, the third of our implicit negative design principles for avoiding unwanted self association was to incorporate structural elements incompatible with beta sheet extension in homo-dimeric species (Fig. 1D). To assess the utility of this principle, we took advantage of the limited number of possible off target edgestrand interactions that can form (Fig. 1C), and docked all protomers against themselves on the edge strand that participates in the heterodimer interface and calculated the Rosetta binding energy after relaxing of the resulting homodimeric dock (Fig. S6A). Homodimer docks of the protomers that chromatographed as monomers in SEC had unfavorable energies compared to those that showed evidence of self association in agreement with our initial hypothesis (Fig. 1D), and visual inspection of these docks suggested that homodimerization was likely prevented by the presence of sterically blocking secondary structure elements (Fig. S6).

In addition to the crystallized fusion proteins (Fig. 2B), 28 more experimentally verified rigid fusion proteins were generated using the 11 base heterodimers and LHD274 (Fig. 3A). The DHR fusions retained both the oligomeric state and binding activity of the unfused counterparts, demonstrating that the designed heterodimers are robust to fusion (Fig. S2E, S5E, S7). With these fusions, there are 74 different possible heterodimeric complexes each with unique molecular scaffolding shapes. The majority of the fusions involve protomers of LHD274 and LHD101. Fusions to LHD101 protomers alone already enable the formation of 30 distinct heterodimeric complexes (Fig. S8).

**Fig. 3.**
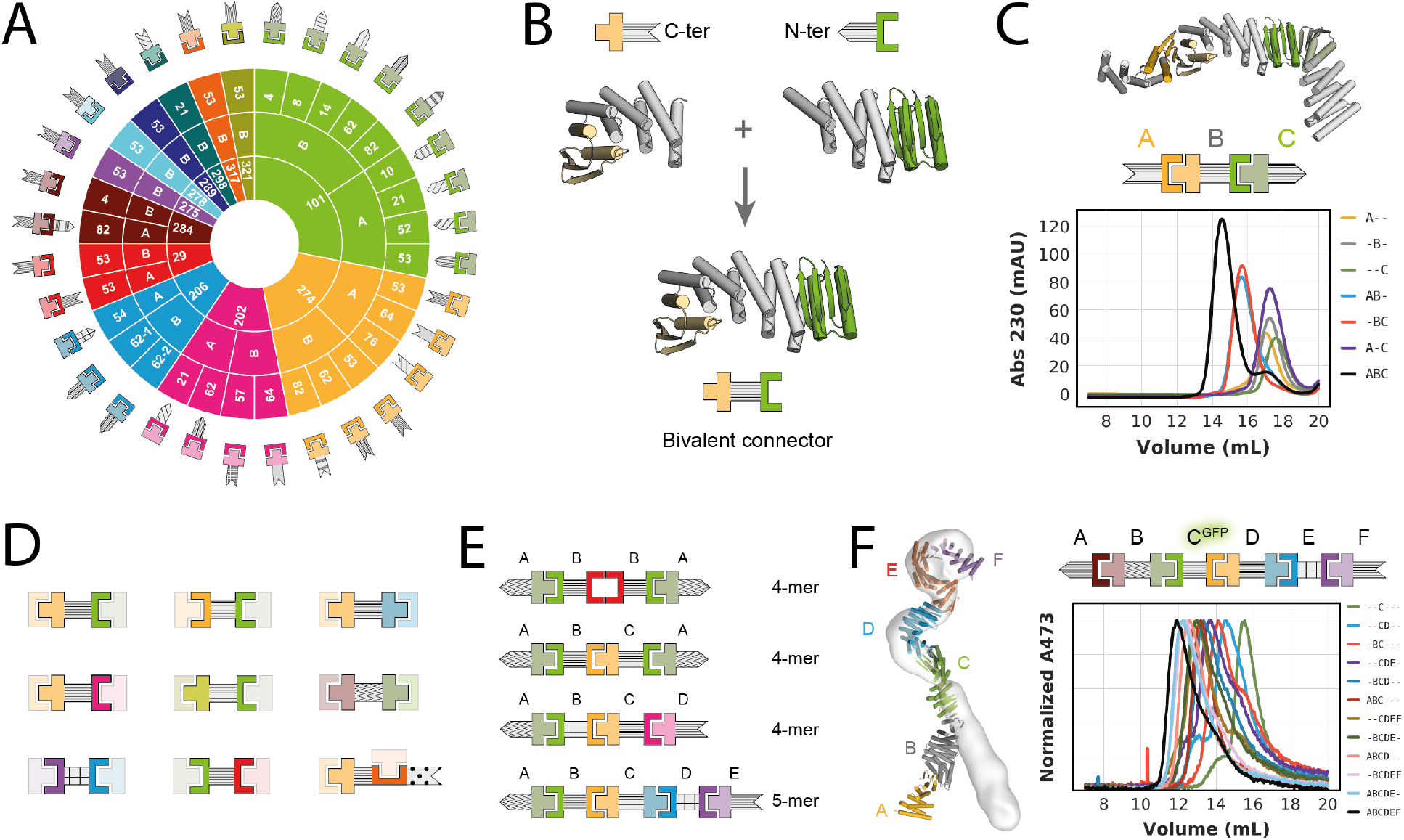
Design of higher order hetero-oligomers. **(A)** Schematic overview of experimentally validated rigid fusion proteins comprising a designed helical repeat protein and a protomer for a heterodimer. **(B)** Schematic representation of the design-free alignment method used to generate bivalent connectors from two of the rigid fusions shown in A. **(C)** Top: Design model and schematic representation of a heterotrimer comprising the bivalent connector shown in B (“B”), and two of the rigid fusions shown in A (“A” and “C”). Bottom: SEC traces for all possible combinations of the trimer components. **(D)** Schematic representations of nine different bivalent connectors that were generated as shown in B and experimentally validated as shown in C (see Fig. S9). **(E)** Schematic representation of experimentally validated higher order assemblies (see Fig. S9 and S10). **(F)** Left: overlay of heterohexamer design model (in colors) and nsEM density (light grey). Right: SEC traces of partial and full mixtures of the hexamer components. Absorbance was monitored at 473 nm to follow the GFP-tagged component C.

Larger multicomponent hetero-oligomeric protein assemblies require subunits that can interact with more than one binding partner at the same time. To this end, we generated single chain bivalent linear connector proteins. We searched for two protomers of different heterodimers that 1) share the same DHR as fusion partner and 2) have compatible termini. Designs fulfilling these criteria can be simply spliced together into a single protein chain on overlapping DHR repeats in a design-free fashion (Fig. 3B). Mixing a linear connector (“B”) with its two cognate binding partners (“A” and “C”) yields a linearly arranged heterotrimer (“ABC”) in which the two terminal capping components A and C are connected through component B, but otherwise are not in direct contact with each other (Fig. 3C). We analyzed the assembly of this heterotrimer and all possible controls by SEC (Fig. 3C), and observed stepwise assembly of the ABC heterotrimer with clear baseline separation from AB and BC heterodimers, as well as from monomeric components (Fig. 3C). Using the 9 different linear connectors created using the above described modular splicing approach (Fig. 3D), we in total assembled 20 heterotrimers including a complex verified by negative-stain electron microscopy (nsEM) (Fig. S9 and S10A).

Linearly arranged hetero-oligomers beyond trimers contain more than one connector subunit in tandem per assembly in contrast to the single connector in heterotrimers. We successfully assembled ABCA and ABCD heterotetramers, each containing two different linear connectors (B and C) and either one or two terminal caps (2xA, or A+D), an ABBA heterotetramer using a homodimeric central connector (2xB) and one terminal cap (2xA), and a negative stain EM verified heteropentamer (ABCDE) containing 3 unique linear connectors and two caps (Fig. 3E, Fig. S9 and S10B). We followed the assembly of an ABCDEF hetero-hexamer in SEC by GFP-tagging one of the components and monitoring GFP absorbance. The full assembly as well as sub-assemblies generated as controls eluted as monodisperse peaks, with elution volumes agreeing well with expected assembly sizes (Fig 3F). Negative stain EM reconstruction of the hexamer confirmed all components were present (Fig. 3F and S10C). Deviation of the experimentally observed shape from the design model likely arises from small inaccuracies in one of the components that cause a lever-arm effect (Fig. 2B).

The design-free generation of bivalent connector proteins from the DHR fusions facilitates the assembly of considerable diversity of asymmetric hetero oligomers. We modularly combined these connectors with each other and with monovalent terminal caps to create 36 hetero-oligomers with up to 6 unique chains which we experimentally validated by SEC and electron microscopy. This number can be readily increased to 489 by including all available components (Fig. 3A,D and supplementary spreadsheet). Since all fusions are rigid helical fusions, the overall molecular shapes of the complexes are well defined allowing control over the spatial arrangement of individual components which could be useful for scaffolding and other applications. Our linear assemblies resemble elongated modular multi-protein complexes found in nature (Fig. S10D), like the Cullin RING E3 Ligases (*28*) that mediate ubiquitin transfer by geometrically orienting the target protein and catalytic domain.

We next sought to go beyond the linear assemblies described thus far and build branched and closed assemblies. Trivalent connectors can be generated from heterodimers in which one protomer has both N- and C-terminal helices (LHD275A, LHD278A, LHD289A, LHD317A). Such protomers can be fused to two helical repeat proteins and spliced together with different halves of other heterodimer protomers via a common DHR repeat (Fig. 3A,B and 4A). The resulting branched connectors (“A”) are capable of binding the three cognate binding partners (“B”,”C”,”D”) simultaneously and conceptually resemble Ste5 and related scaffolding proteins that organize MAP kinase signal transduction pathways in eukaryotes (*29*). Through SEC analyses we verified the assembly of two different tetrameric branched ABCD complexes, each containing one trivalent branched connector bound to three terminal caps (Fig. 4B and S11A,B). For one of these, the complex was confirmed by negative stain EM class averages and 3D reconstructions indicating not only that all binding partners are present, but also that the shape closely matches the designed model (Fig. 4A and S11A).

**Fig. 4.**
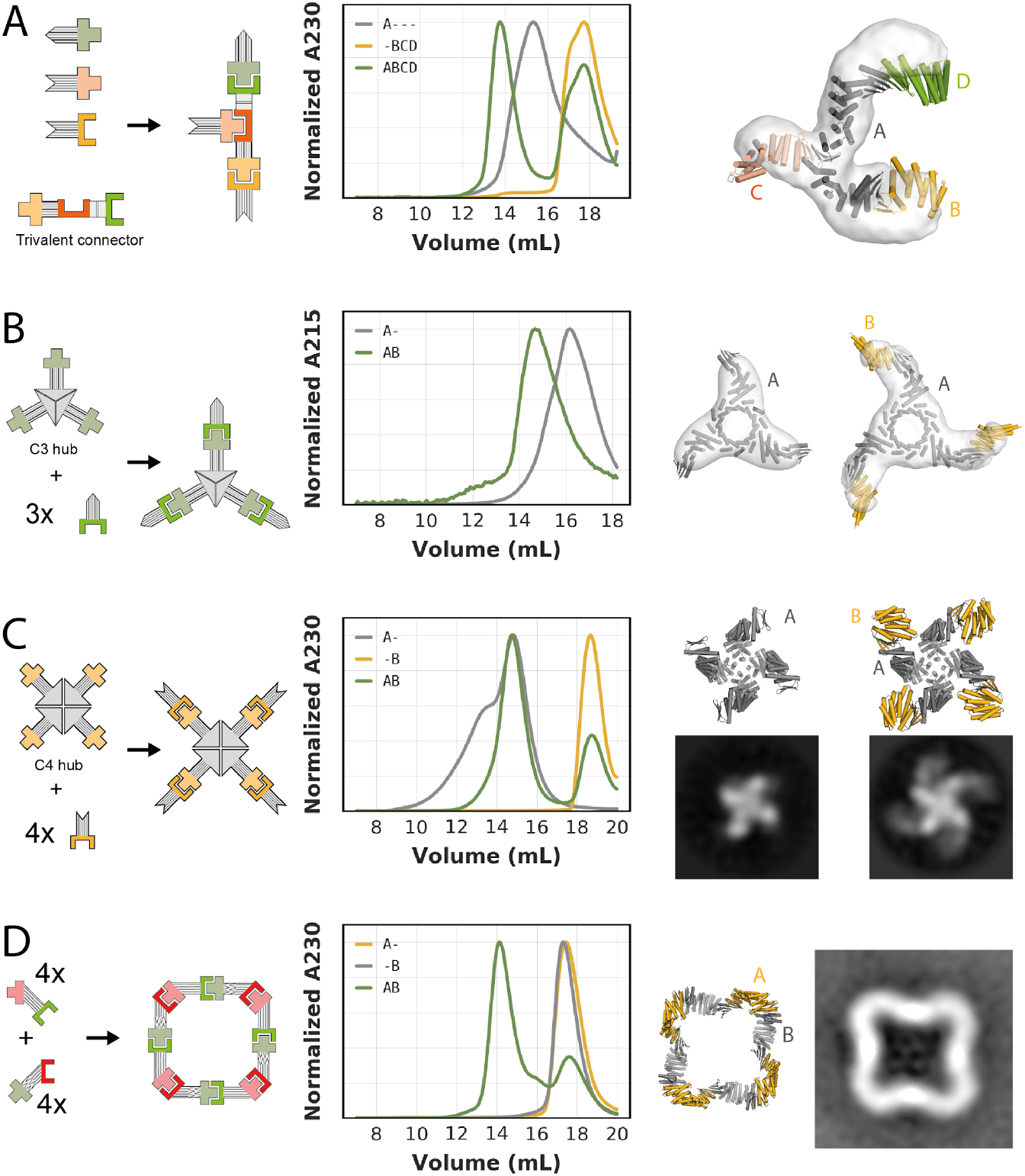
Design of branched and closed hetero-oligomeric assemblies. **(A)** Left: Schematic representation of a trivalent connector (“A”) that can bind three different binding partners (“B”, “C”, “D”). Center: SEC analysis of the trivalent connector, the binding partners, and the full assembly mixture. Right: Overlay of design model (in colors) and nsEM density (light grey) of the complex formed by the trivalent connector and all three binding partners. **(B)** From left to right: : Schematic representation of a C3-symmetric “hub” that can bind three copies of one binding partner; SEC analysis of the C3-symmetric “hub” without (“A-”) and with (“AB”) binding partner; overlay of design model (dark grey) and nsEM density (light grey) of the C3-symmetric “hub”; overlay of design model (dark grey and gold) and nsEM density (light grey) of the C3-symmetric “hub” bound to three copies of its binding partner. **(C):** From left to right: : Schematic representation of a C4-symmetric “hub” that can bind four copies of one binding partner; SEC analysis of the C4-symmetric “hub” without (“A-”) and with (“AB”) binding partner; design model (top) and representative nsEM class average (bottom) of the C4-symmetric “hub”; design model (top) and representative nsEM class average (bottom) of the C4-symmetric “hub” bound to 4 copies of the binding partner. **(D)** From left to right: : Schematic representation of a C4-symmetric closed ring comprising two components (“A” and “B”); SEC analysis of the individual ring components (“A-” and “-B”) and the stoichiometric mixture (“AB”); design model of the C4-symmetric ring; representative nsEM class average.

A different type of branched assemblies are “star shaped” oligomers with cyclic symmetries, akin to natural assemblies formed by IgM and the Inflammasome (*30, 31*). Using the design-free alignment approach described above (Fig. 3B), we fused our new building blocks (Fig. 3A) to previously designed homo-oligomers (*22, 32*), that terminate in helical repeat proteins (Fig. 4B,C). Such fusions yield central homo oligomeric hubs (“A_n”) that can bind multiple copies of the same binding partner (“n*B”). We generated C3- and C4-symmetric “hubs” that can bind 3 or 4 copies of their binding partners, respectively (Fig. 4B,C). In both cases, the oligomeric hubs are stable and soluble in isolation and readily form the target complexes when mixed with their binding partners, as confirmed by SEC chromatography, negative stain EM class averages and 3D reconstructions (Fig. 4B,C and Fig. S11C, S12). For the C4-symmetric hub in the absence of its binding partner we observed an additional concentration-dependent peak on SEC (Fig. 4C, Fig. S12A), indicating formation of a higher-order complex. This is likely a dimer of C4 hubs, since the C4 hub contains the redesigned protomer LHD274B, that despite its reduced homodimerization propensity compared to parent design LHD29B still weakly homodimerizes (Fig. S4). Notably, addition of the binding partner disrupted the higher order assembly, yielding the on-target octameric (A4B4) complex (Fig. 4C), illustrating this system can reconfigure.

In addition to linear and branched assemblies, we designed closed symmetric two-component assemblies. Designing these presents a more complex geometric challenge, as the interaction geometry of all pairs of subunits must be compatible with a single closed three dimensional structure of the entire assembly. We used architecture-aware rigid helical fusion (*7, 33*) to generate two bivalent connector proteins from the crystal-verified fusions of LHD29 and LD101 (Fig 2B) that allow assembly of a perfectly closed C4-symmetric hetero-oligomeric two-component ring (Fig. 4D). Individually expressed and purified components are stable and soluble monomers in isolation, as confirmed by SEC and native MS (Fig. 4D, Fig. S13). Upon mixing, the components form a higher-order complex that by native MS comprises four copies of each component. Negative stain EM confirms that this higher-order complex is nearly identical to the designed C4 symmetric ring (Fig. 4D, Fig. S13). Using our heterodimeric building blocks, the same architecture-aware fusion method could potentially be used to design a variety of different closed symmetric complexes that assemble from well-behaved components.

Because our designed building blocks are stable in solution and not kinetically trapped in off-target homo-oligomeric states, the assemblies they form can rapidly reconfigure, as outlined in Figure 1A and observed for the C4-symmetric hub shown in Figure 4C. We further evaluated this reconfigurability using two different approaches to assemble and then reconfigure a heterotrimer. First, we assembled an ABC trimer using a GFP-tagged version of a linear connector B and unfused terminal caps A and B (Fig. 5A). The pre-incubated trimer was next mixed with either buffer or a DHR fusion variant of component C, called C’. As indicated by the shift of the trimer peak in SEC, component C (8.6 kDa) readily exchanged with C’ (27.7 kDa), to form a larger ABC’ complex. Subunit exchange was confirmed by biolayer interferometry (Fig. S14).

**Fig. 5.**
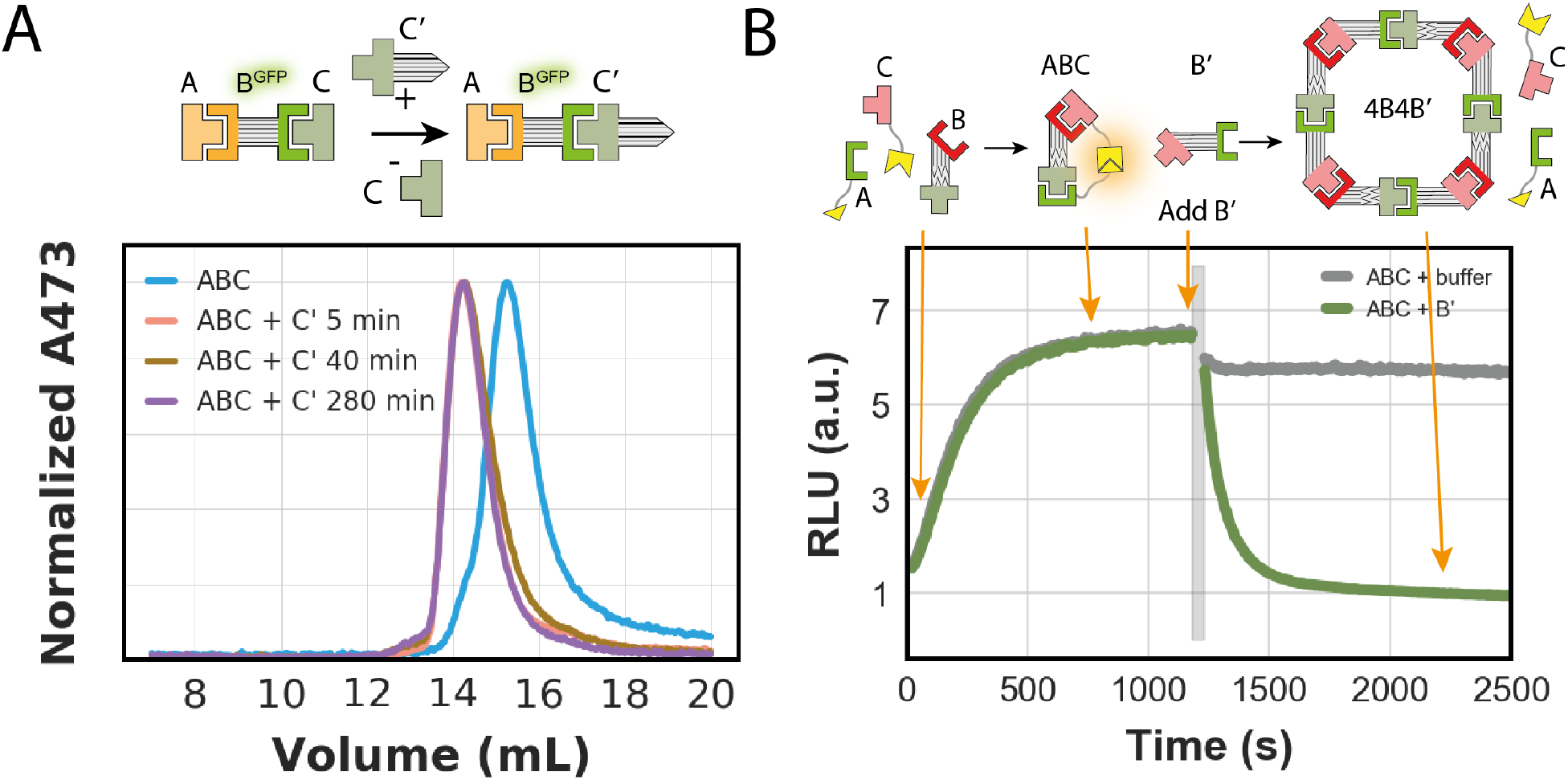
Dynamically reconfigurable protein assemblies. **(A)** Exchange experiment in which a pre-assembled trimer (“ABC”) is incubated with a variant of one of the components (“ C’ “). Top: Schematic representation, bottom: SEC traces of trimer mixture before and after addition of component C’. **(B)** Top: schematic representation of a split luciferase experiment in which two protomers (“A” and “C”) are fused to split luciferase parts (yellow shapes). Bottom: Real-time luminescence measurement of two samples containing the mixture “ABC” shown on the left. Grey bar indicates addition of either buffer (grey trace) or component B’ (green trace).

Second, we followed the transition, through subunit exchange, of a linear heterotrimer to the designed C4 symmetric hetero-oligomeric two-component ring using an *in vitro* split luciferase reporter assay (Fig. 5B). We first assembled an ABC heterotrimer, in which chain B is one of the two components of the ring, and A and C are the corresponding terminal cap binding partners fused to the two parts of the split luciferase. In absence of B, components A and C do not interact. Upon addition of B, the heterotrimer forms, resulting in luciferase activity. Subsequent addition of the second component of the C4 symmetric ring, B’, led to a rapid decrease in luciferase activity, indicating disassembly of the trimer (Fig. 5B) consistent with ring formation from the two components observed in SEC (Fig. 4C). Taken together, these experiments indicate that subunit exchange can take place on the several minute time scale and pave the way for applications that require designed dynamic reconfigurability of multiprotein complexes.

## Conclusion

Our implicit negative design principles enable the de novo design of heterodimer pairs for which the individual protomers are stable in solution and readily form their target heterodimeric complexes upon mixing. Rigid fusion of multiple halves of heterodimers onto DHR proteins enables the design of higher order asymmetric multiprotein complexes that range in shape from linear and cyclic to branched. The large number of characterized rigid fusions with different shapes and the modular nature of our assembly platform enables fine tuning of protein complex geometries, for example by changing the number of repeats in the DHR proteins and using the same heterodimer half fused to different DHRs.

Since the unfused protomers are small (between 7 and 15 kDa without DHR or tags), they can be readily fused to target proteins of interest. Our bivalent or trivalent connectors can then be used to colocalize and geometrically position two or three such target protein fusions, respectively, and our symmetric hubs can be used to colocalize and position multiple copies of the same target fusion. Due to the modularity of our system, the same set of target fusions can be arranged in multiple different arrangements with adjustable distances, angles, and copy numbers by simply using different connectors. Since all components are soluble and well-behaved in isolation, stepwise assembly schemes are possible in which, for example, two constitutively expressed target protein fusions do not interact until expression of a connector is induced, leading to formation of a trimeric complex. Using one of our ABCD tetramers, such a system could be extended to enable simple logic operations: two target proteins fused to components A and D will only be colocalized if both B and C are present. Since the thermodynamic and kinetic properties of our heterodimers are not altered by rigid fusions, the behaviour of multi-component assemblies can be predicted based on the properties of the individual interfaces (compare Fig. S5F,G). Like the sophisticated multi-protein complexes that govern cellular processes in biology (and unlike most previously designed, highly symmetric protein assemblies) our designed assemblies can reconfigure by addition of new subunits and loss of already incorporated ones, opening the door to a wide range of new applications for de novo protein design.

## Supporting information

supplementary_pdbs_scripts

## Acknowledgments

We acknowledge Baker lab members for discussion, beamline scientists at the APS for crystallography support, Florian Busch and Andrew Norris for native mass spectrometry measurements, and Titia Sixma and Hendrik Dietz for critical reading of the manuscript.

## Funding

EMBO long term fellowship ALTF 1295-2015 and ALTF 139-2018 (DDS, BIMW)

Washington Research Foundation Innovation Fellowship (DDS)

Human Frontiers Science Program long term fellowship (FP, AC)

DARPA Biostasis HR001118S0034 (YH, DB)

Open Philanthropy Project Improving Protein Design Fund (DB, FP, AKB, AK),

The Audacious Project at the Institute for Protein Design (LMM, HMM, BJRT, NIE)

Eric and Wendy Schmidt by recommendation of the Schmidt Futures (DB, YH)

The Howard Hughes Medical Research Institute (DB, AC).

NIH Resource for Native Mass Spectrometry Guided Structural Biology P41 GM128577 (Vicki Wysocki, Ohio State University)

## Author contributions

D.D.S. and F.P. developed hetero-oligomer design pipeline, performed design calculations and experiments and analyzed all data. A.C. designed and characterized homo-oligomeric C3 hub. Y.H. designed and characterized two component C4 ring. B.I.M.W. performed and analyzed split luciferase binding assays. N.I.E. designed and characterized homo-oligomeric C4 hub. A.C., Y.H. and N.I.E. performed negative stain EM and 3D reconstructions. B.J.R.T. designed scaffolds. L.M.M. and H.M.M. purified designs. D.D.S., A.B. and A.K. determined crystal structures.

## Competing interests

DDS, FP, AC, NIE, YH, BJRT and DB are inventors on a provisional patent application submitted by the University of Washington for the design, composition and function of the proteins created in this study.

## Data and materials availability

Crystallographic models have been deposited in the RCSB PDB (accession codes: 6wmk, 7mwq, 7mwr) and will be released upon publication. All data are available in the main text or the supplementary materials.

## Supplementary Materials

### Materials and Methods

Figs. S1 to S14

Tables S1 to S4

References (*34–55*)

Data S1_components_and_assemblies.xlsx to S2_pdbs_and_scripts.tar.gz

## Supplementary Materials for

**Other Supplementary Materials for this manuscript include the following:**

Data S1 to S2: S1_components_and_assemblies.xlsx, S2_pdbs_and_scripts.gz

### Materials and Methods

#### Protein design

##### Docking procedure

As scaffolds for generating edge-strand heterodimers we used mixed alpha/beta proteins designed by citizen scientist (*21*) and variants of the fold-it scaffolds that were either expanded with additional helices (see backbone generation methods), and/or fused to de novo helical repeat (DHR) proteins (*27*). Edgestrand docking was performed as described previously (*18*). Exposed edgestrands suitable for docking were identified by calculating the solvent accessible surface area of beta sheet backbone atoms in all the scaffolds used in the docking procedure. Next, the c-alpha atoms of each strand of short 2 stranded parallel and antiparallel beta sheet motifs were aligned to the exposed edge strand yielding an aligned clashing strand and free dock strand. After removal after the aligned clashing strand, the docked strand was trimmed at N and/or C terminus in order to remove potential clashes and subsequently minimized using Rosetta FastRelax (*34*) to optimize backbone to backbone hydrogen bonds. Docks failing a specified threshold value (typically -4 using ref2015) for the backbone hydrogen bond scoreterm in Rosetta (hbond_lr_bb) were discarded. The minimized docked strands were next geometrically matched to the scaffold library using the MotifGraftMover to create a docked protein-protein complex (*35*).

##### Interface design

The interface residues of the docked heterodimer complexes were optimized using Rosetta combinatorial sequence (*36–39*) design using “ref2015”,”beta_nov16” or “beta_genpot” as scorefunctions (*40*). The interface polarity of the docked heterodimer complexes were fine tuned in several ways (see supplement for description of design xml’s). First, the HBNetMover (*11*) was used to install explicit hydrogen bond networks containing at least 3 hydrogen bonds across the interface. Later design rounds consisted of two seperate interface sequence optimization steps. First interface residues were optimized without compositional constraints yielding a substantial number of hydrophobic interactions in the interface. The best designs were subsequently selected and hydrophobic residue pairs with the lowest Rosetta energy interactions across the interface were stored as a seed hydrophobic interaction hotspot. In a second round, a polar interaction network was designed around the fixed hydrophobic hotspot interaction using compositional constraints that favor polar interactions (*26*). Designs were filtered on interface properties such as binding energy, buried surface area, shape complementarity, degree of packing, and presence of unsatisfied buried polar atoms. A final selection was made by visual inspection of models.

##### Backbone generation and scaffold design

*De novo* designed protein scaffolds created by fold-it players (*21*) were expanded with C-terminal polyvaline helices using blueprint based backbone generation (*23, 24*). The amino acid identities of the newly built helices and their surrounding region were optimized using Rosetta combinatorial sequence designs using a flexible backbone. The resulting models were folded *in silico* using Rosetta folding simulations and trajectories that converged to the designed model structure without off-target minima were selected for rigid fusion and heterodimer design.

##### Design of rigid fusions

To generate rigid fusions of scaffolds or heterodimers to DHRs we adapted the HFuse pipeline (*22*), (*7*): Fusion junctions were designed using the Fastdesign mover allowing backbone movement, and additional filters were included to ensure sufficient contact between DHR and scaffold/heterodimer. When fusing to heterodimers, an additional filter was employed to prevent additional contacts between the DHR and the other protomer of the dimer. Bivalent connectors were generated by aligning two proteins that share the same DHR along their shared helical repeats, and subsequently splicing together the sequences. To build the C3-symmetric “hub”, we used a previously published 12x toroid crystal structure (*32*). The starting structure was relaxed, Z axis aligned, and cut into three C3 symmetric chains. Then the HFuse software (*22*), (*7*) was used to sample DHR fusions to the exposed helical C-termini, and the newly created interfaces were redesigned using RosettaScripts. For the C4 symmetric hub, we used a previously published C4-symmetric homooligomer that already containe a n-terminal DHR. For both hubs, matching DHR fusions of heterodimer protomers we then used the same align and splice approach as for the bivalent connectors.

##### Design of C4 rings

Using the relaxed crystal structures of LHD29 and LHD101 fused to their respective DHRs, the WORMS software (*7, 9, 33*) was used to fuse the two hetero-dimers into cyclic symmetrical rings. As one construct has exposed N-termini and the other has exposed C-termini, they were able to be fused head to tail without introduction of further building blocks. Briefly, the first 3 repeats of each repeat protein was allowed to be sampled as fusion points to ensure that the heterodimer interface was not altered. Following fusion into cyclic structures, fixed backbone junction design was applied to the new fusion point using RosettaScripts (*38*), optimizing for shape complementarity (*41*). One design from each symmetry: C3, C4, C5, and C6 were selected for experimental testing.

##### Protein expression and purification

Synthetic genes encoding designed proteins and their variants were purchased from Genscript or Integrated DNA technologies (IDT). Bicistronic genes were ordered in pET29b with the first cistron being either without tag or with an N-terminal sfGFP tag followed by the intercistronic sequence TAAAGAAGGAGATATCATATG. The second cistron was tagged with a polyhistidine His6x tag at the C-terminus. Plasmids encoding the individual protomers were ordered in pET29b either with or without Avi-Tag, with an N-terminal polyhistidine His6x tag followed by a TEV cleavage site, N-terminal polyhistidine His6x tag followed by a snac cleavage site or C-terminal polyhistidine His6x tag preceded by a snac tag (see supplementary spreadsheet for detailed construct information). Proteins were expressed in BL21 LEMO E.coli cells by autoinduction using TBII media (Mpbio) supplemented with 50×5052, 20 mM MgSO4 and trace metal mix, or in almost TB media containing 12 g peptone and 24 g yeast extract per liter supplement with 50×5052, 20 mM MgSO4, trace metal mix and 10x phosphate buffer. Proteins were expressed under antibiotics selection at 37 degrees overnight or at 18 degrees for 24h after initial growth for 6-8h at 37 degrees. Cells were harvested by centrifugation at 4000x g and lysed by sonication after resuspension of the cells in lysis buffer (100 mM Tris pH 8.0, 200 mM NaCl, 50 mM Imidazole pH 8.0) containing protease inhibitors (Thermo Scientific) and Bovine pancreas DNaseI (Sigma-Aldrich). Proteins were purified by Immobilized Metal Affinity Chromatography. Cleared lysates were incubated with 2-4ml nickel NTA beads (Qiagen) for 20-40 minutes before washing beads with 5-10 column volumes of lysis buffer, 5-10 column volumes of high salt buffer (10 mM Tris pH 8.0, 1 M NaCl) and 5-10 column volumes of lysis buffer. Proteins were eluted with 10 ml of elution buffer (20 mM Tris pH 8.0, 100 mM NaCl, 500 mM Imidazole pH 8.0).

Designs were finally polished using size exclusion chromatography (SEC) on either Superdex 200 Increase 10/300GL or Superdex 75 Increase 10/300GL columns (GE Healthcare) using 20 mM Tris pH 8.0, 100 mM NaCl or 20 mM Tris pH 8.0, 300 mM NaCl. Cyclic assemblies of C3 and C4 symmetries were purified using a Superose 6 increase 10/300GL (GE Healthcare). The two component C4 rings were SEC purified in 25 mM Tris pH 8.0, 300 mM NaCl. Peak fractions were verified by SDS-PAGE and LC/MS and stored at concentrations between 0.5-10 mg/ml at 4 degrees or flash frozen in liquid nitrogen for storage at -80. Designs that precipitated at low concentration upon storage at 4 degrees could in general be salvaged by increasing the salt concentration to 300-500 mM NaCl.

For structural studies, designs with a polyhistidine tag and TEV recognition site were cleaved using TEV protease (his6-TEV). TEV cleavage was performed in a buffer containing 20 mM Tris pH 8.0, 100 mM NaCl and 1 mM TCEP using 1% (w/w) his6-TEV and allowed to proceed o/n at room temperature. Uncleaved protein and his6-TEV were separated from cleaved protein using IMAC followed by SEC. Designs carrying a C-terminal SNAC-polyhistine tag (GGSHHWGS(…)HHHHHH) were cleaved chemically via on-bead nickel assisted cleavage (*42*): nickel bound designs were washed with 10CV of lysis buffer followed by 5CV of 20 mM Tris pH 8.0, 100 mM NaCl. Proteins were subsequently washed with 5CV of SNAC buffer (100 mM CHES, 100 mM Acetone oxime, 100 mM NaCl, pH 8.6). Beads were next incubated with 5CV SNAC buffer + 2 mM NiCl_2_ for more than 12 hours at room temperature on a shaking platform to allow cleavage to take place. Next, the flow through containing cleaved protein was collected. The flow throughs of two additional washes (SNAC buffer/SNACbuffer+50 mM Imidazole) of 3-5CV were also collected to harvest any remaining weakly bound protein. Cleaved proteins were finally purified by SEC.

##### Luciferase binding assays

Assays were performed in 20 mM sodium phosphate, 100 mM NaCl, pH 7.4, 0.05% v/v Tween 20. Reactions were assembled in 96 well plates (Corning, cat #3686) in the presence of Nano-Glo substrate (Promega, cat. #N1130) diluted 100x or 500x for kinetics and endpoint measurements respectively (see supplement for detailed information). Luminescence was recorded on a Synergy Neo2 plate reader (BioTek). Kinetic assays were performed under pseudo first-order conditions, with the final concentration of one protein at 1 nM and the other at 10 nM. Dead times between substrate addition and data acquisition were typically 15-30s. For long kinetic measurements (Fig. S5A), mastermixes of the protein complexes were made and aliquots were sampled at regular intervals. Data were fitted (*43*) to a single exponential decay function:

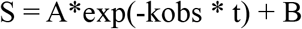

where *t* is time, *S* is the luminescence signal, and the fitted parameters are: *A* the amplitude, *k*_obs_ the observed rate constant, and *B* the endpoint luminescence.

Equilibrium binding reactions were incubated overnight at room temperature before adding substrate and immediately measuring luminescence. The data was fitted to the following equation to obtain *K*_d_ values:

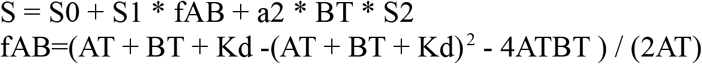

where *A_T_* and *B*_T_ are the total concentrations of each species ( *A_T_* = 1 nM, *B_T_* is the titrated species), and *S* is the observed signal. The fitted parameters are: *S_0_* the pre-saturation baseline, *S_1_* the post-saturation baseline, *a_2_* and *S_2_* the correction terms, and *K_d_* the equilibrium dissociation constant.

ABC complex equilibrium binding experiments were performed using the concentration indicated in the figure legend of Fig. S5G for the constant components, and titrating B. Reactions were incubated overnight before adding substrate and data acquisition (for details on the modeling of ABC kinetics see supplement). For the ABC reconfiguration kinetics (Fig. 5B) components A (0.5 nM) and C (200 nM) were briefly pre-incubated in the presence of substrate, before adding component B (50 nM) to start the reaction. At equilibrium component B’ (2000 nM) was added to the reactions, and data acquisition was resumed until dissociation was complete.

##### Enzymatic protein biotinylation

Avi-tagged (GLNDIFEAQKIEWHE, see supplement) proteins were purified as described above. The BirA500 (Avidity, LLC) biotinylation kit was used to biotinylate 840 uL of protein from the IMAC elution in a 1200 uL (final volume) reaction according to the manufacturer’ protocol. Reactions were incubated at 4 degrees C o/n and purified using size exclusion chromatography on a Superdex 200 10/300 Increase GL (GE Healthcare) or S7510/300 Increase GL (GE Healthcare) in SEC buffer (20 mM Tris pH 8.0, 100 mM NaCl).

##### Biolayer interferometry

Biolayer interferometry experiments were performed on an OctetRED96 BLI system (ForteBio, Menlo Park, CA). Streptavidin coated biosensors were first equilibrated for at least 10 minutes in Octet buffer (10 mM HEPES pH 7.4, 150 mM NaCl, 3 mM EDTA, 0.05% Surfactant P20) supplemented with 1 mg/ml Bovine Serum Albumin (SigmaAldrich). Enzymatically biotinylated designs were immobilized onto the biosensors by dipping the biosensors into a solution with 10-50 nM protein for 30-120 s. This was followed by dipping in fresh octet buffer to establish a baseline for 120 s. Titration experiments were performed at 25 °C while rotating at 1,000 r.p.m. Association of designs was allowed by dipping biosensors in solutions containing designed protein diluted in octet buffer until equilibrium was approached followed by dissociation by dipping the biosensors into fresh buffer solution in order to monitor the dissociation kinetics. Steady-state and global kinetic fits were performed using the manufacturer’s software (Data Analysis 9.1) assuming a 1:1 binding model.

##### SEC binding assays

Complexes and individual components were diluted in 20 mM Tris pH 8.0, 100 mM NaCl. After o/n equilibration of the mixtures at room temperature or 4 degrees C, 500 ul of sample was injected onto a Superdex 200 10/300 increase GL (dimers, linear assemblies) or Superose 6 increase 10/300 GL (symmetric assemblies) (all columns from GE healthcare) using the absorbance at 230 nm or 473 nm (for GFP tagged components) as read-out. Dimers were mixed at monomer concentrations of 5 µM or higher. Trimer and ABCD tetramer mixtures contained 5 µM of the bivalent connector, and 7.5 µM of each terminal cap (lower absolute concentrations with the same ratios were used for some trimers). ABCA tetramer mixtures contained 5 µM per bivalent connector and 15 µM terminal cap. The hexamer mixture contained 3 µM of components C and D, 3.6 µM of B and E, and 4.4 µM of A and F. The branched assembly shown in Figure 4A contained 2.8 µM of the trivalent connector and 4 µM of each cap. For the exchange experiment shown in Fig. 5A, the ABC trimer was preincubated at concentrations of 6 µM B and 9 µM each of A and C. C’ was then added to reach a final concentration of 2 µM B, 3 µM each of A and C, and 6 µM C’.

Native mass spectrometry

Sample purity, integrity, and oligomeric state was analyzed by on-line buffer exchange MS in 200 mM ammonium acetate using a Vanquish ultra-high performance liquid chromatography system coupled to a Q Exactive ultra-high mass range Orbitrap mass spectrometer (Thermo Fisher Scientific). A self-packed buffer exchange column was used (P6 polyacrylamide gel, BioRad) (*44*). The recorded mass spectra were deconvolved with UniDec version 4.2+ (*45*).

##### Crystal structure determination

For all structures, starting phases were obtained by molecular replacement using Phaser (*46*). Diffraction images were integrated using XDS (*47*) or HKL2000 (*48*) and merged/scaled using Aimless (*49*). Structures were refined in Phenix (*50*) using phenix.autobuild and phenix.refine or Refmac (*51*). Model building was performed using COOT (*52*).

Proteins were crystallized using the vapor diffusion method at room temperature. LHD29 crystals grew in 0.2M Sodium Iodide, 20% PEG3350, LHD29A53/B53 crystals in E5 and LHD101A53/B4 crystals in 2.4M Sodium Malonate pH 7.0. Crystals were harvested and cryoprotected using 20% PEG200 for LHD29, 20% PEG400 for LHD29A53/B53 and 20% glycerol for LHD101A53/B4 before data was collected at the Advanced Light Source (Berkeley, USA). The structures were solved by molecular replacement using either computationally designed models of individual chains A or B or the full heterodimer complex as search models.

##### Electron microscopy

SEC peak fractions were concentrated prior to negative stain EM screening. Samples were then immediately diluted 5 to 150 times in TBS buffer (25 mM Tris pH 8.0, 25 mM NaCl) depending on sample concentration. A final volume of 5 μL was applied to negatively glow discharged, carbon-coated 400-mesh copper grids (01844-F, TedPella,Inc.), then washed with Milli-Q Water and stained using 0.75% uranyl formate as previously described (*53*). Air-dried grids were imaged on a FEI Talos L120C TEM (FEI Thermo Scientific, Hillsboro, OR) equipped with a 4K × 4K Gatan OneView camera at a magnification of 57,000x and pixel size of 2.51. Micrographs were imported into CisTEM software or cryoSPARC software and a circular blob picker was used to select particles which were then subjected to 2D classification. Ab initio reconstruction and homogeneous refinement in Cn symmetry were used to generate 3D electron density maps (*54, 55*).

#### Additional methods for the Luciferase assay

##### Constructs

Split luciferase reporter constructs were ordered as synthetic genes from Genscript. Each design was N-terminally fused to a sfGFP (for protein quantification in lysate), and C-terminally fused to either smBiT or lgBiT of the split luciferase components. A Strep-tag was included at the N-terminus for purification, and a GS-linker was inserted between the design and the split luciferase component.

##### Expression for multiplexed assay

Plasmids were transformed into Lemo21(DE3) cells (New England Biolabs), and grown in 96 deepwell plates overnight at 37 °C in 1 mL of LB containing 50 ug/mL of kanamycin sulfate. The next day, 100 uL of overnight cultures were used to inoculate 96 deepwell plates containing 900 uL of TBII medium (MP Biomedicals) with 50 ug/mL of kanamycin sulfate, and the cultures were grown for 2 h at 37 °C before induction with 0.1 mM IPTG. Protein expression was carried out at 37 °C for 4 h before the cells were harvested by centrifugation (4,000 x g, 5 min). Cell pellets were resuspended in 100 uL of lysis buffer (10 mM sodium phosphate, 150 mM NaCl, pH 7.4, 1 mg/mL lysozyme, 0.1 mg/mL DNAse I, 5 mM MgCl_2_, 1 tablet/50 mL of cOmplete protease inhibitor (Roche), 0.05% v/v Tween 20), and cell were lysed by performing three freeze/thaw cyles (1 h incubations at 37 °C followed by freezing at -80 °C). The lysate was cleared by centrifugation (4,000 x g, 20 min), and the soluble fraction transferred to a 96 well assay plate (Corning, cat #3991). Concentrations of the constructs in soluble lysate were determined by sfGFP fluorescence using a calibration curve.

##### Lysate production for multiplexed assay

Neutral lysate for preparing serial dilutions was prepared by transforming Lemo21(DE3) with the pUC19 plasmid. Transformations were used to inoculate small overnight cultures, which were used to inoculate 0.5 L TBII cultures (all cultures contained 50 ug/mL of carbenicillin). Cells were grown for 24 h at 37 °C before being harvested. Pellets were resuspended in the same lysis buffer, followed by sonication. The lysate density was adjusted with lysis buffer to have its OD280 matching pUC19 control wells from the 96 well expression plate.

##### Expression and purification

Plasmids were transformed into Lemo21 (DE3) cells, and used directly to inoculate 50 mL of auto-induction media (TBII supplemented with 0.5 % w/v glucose, 0.05% w/v glycerol, 0.2% w/v lactose monohydrate, and 2 mM MgSO_4_, 50 ug/mL kanamycin sulfate). The cultures were incubated at 37 °C for 20-24 h, before harvesting the cells by centrifugation (4,000 x g, 5 min). Cells were resuspended in 10 mL of lysis buffer (100 mM Tris, 150 mM NaCl, pH 8, 0.1 mg/mL lysozyme, 0.01 mg/mL DNAse I, 1 mM PMSF) and lysed by sonication. The insoluble fraction was cleared by centrifugation (16,000 x g for 45 min), and the proteins were purified from the soluble fraction by affinity chromatography using Strep-Tactin XT Superflow High-Capacity resin (IBA Lifesciences). Elutions were performed with 100 mM Tris, 150 mM NaCl, 50 mM biotin, pH 8, and the proteins were further purified by size-exclusion chromatography using a Superdex 200 10/300 increase column equilibrated with 20 mM sodium phosphate, 100 mM NaCl, pH 7.4, 0.05% v/v Tween 20.

##### Binding assays

All assays were performed in 20 mM sodium phosphate, 100 mM NaCl, pH 7.4, 0.05% v/v Tween 20. Depending on the source of the protein used in the assay (purified components or lysate), soluble lysate components were also present. Reactions were assembled in 96 well plates (Corning, cat #3686) in the presence of Nano-Glo substrate (Promega, cat. #N1130) diluted 100x or 500x for kinetics and endpoint measurements respectively, and the luminescence signal was recorded on a Synergy Neo2 plate reader (BioTek).

Kinetic binding assays were performed under pseudo first-order condtions, with the final concentration of one protein at 1 nM and the other at 10 nM. Stock solutions were mixed in a 1:1 volume ratio in the presence of substrate, and the dead-time between mixing and starting the measurement (typically 15-30 s) added during data-processing. For long kinetic measurements (Fig. S5A), the proteins were pre-mixed, and kept in a sealed tube at room temperature over the course of the experiment. Aliquots were taken at regular intervals, mixed with substrate, and immediately recorded. All kinetic measurements were fitted to a single exponential decay function:

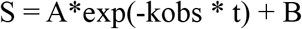

where *t* is time (the independent variable), *S* is the observed luminescence signal (the dependent variable), and the fitted parameters are: *A* the amplitude, *k*_obs_ the observed rate constant, and *B* the endpoint luminescence.

Equilibrium binding assays were performed with one component kept constant at 1 nM while titrating the other protein. Serial dilutions curves were prepared over 12 points, with a ¼ dilution factor between each step. The concentration of protein in the soluble lysate provided the highest concentration point of the curve. To avoid serial dilution of the other lysate components, all stocks were prepared with neutral lysate. The assembled plates were incubated overnight at room temperature before adding substrate and immmediately measuring luminescence. The data was fitted to the following equation to obtain *K*_d_ values:

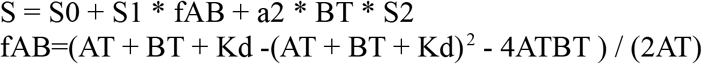

where *A_T_* and *B*_T_ are the total concentrations of each species (the independent variables, *A_T_* = 1 nM, *B_T_* is the titrated species), and *S* is the observed signal (the dependent variable). The fitted parameters are: *S_0_* the pre-saturation baseline, *S_1_* the post-saturation baseline, *a_2_* and *S_2_* the correction terms, and *K_d_* the equilibrium dissociation constant.

Specificity matrices were obtained by preparing all combinations of smBiT and lgBiT proteins at 100 nM and 1 nM final concentrations respectively. The reactions were incubated overnight at room temperature before adding substrate and immediately measuring luminescence.

Ternary complex equilibrium binding experiments were performed with pure protein, using the concentration indicated in the figure legend of Fig. S5G for the constant components, and titratring B. After assembly, the plates were incubated overnight before adding substrate and immediately measuring luminescence.

Ternary complex reconfiguration kinetics (Fig. 5B) were measured with pure proteins. Components A and C were briefly pre-incubated in the presence of substrate, before adding component B to start the reaction. Once the association was complete, the assay plate was briefly taken out of the plate reader, component B’ was added to the reactions, and data acquisition was resumed until dissociation was complete.

##### Simulation of ternary complex

Systems of ordinary differential equations describing the kinetics of interactions between the species involved in the formation of the ternary complex were numerically integrated using integrate.odeint() as implemented in Scipy (version 1.6.3). Steady-state values were used to determine the distribution of species at thermodynamic equilibrium.

The ternary system is composed of the following species: A, B, C, AB, BC, ABC. The following set of equations was used to describe the system:

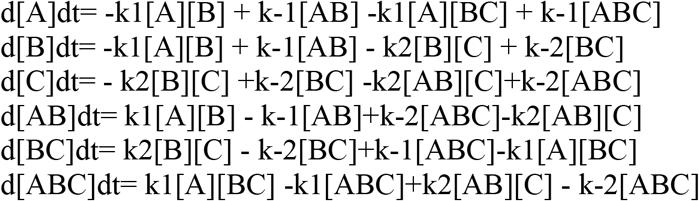

where ki describe bimolecular association rate constants and k-irepresent unimolecular dissociation rate constants. K1=k-1/ k1, and K2=k-2 / k2describe the affinity of the A:B and B:C interfaces respectively.

Data were collected from one crystal per condition. ^a^ Values given in parentheses refer to reflections in the outer resolution shell. For calculation of *R*_free_, 5% of all reflections were omitted from refinement.

#### Data S1. S1_components_and_assemblies.xlsx

Spreadsheet with sequences and parameters for proteins and assemblies shown in this work. Tab1: LHD components. Sequences and parameters for all heterodimers, fusions, connectors, and hubs presented in this work. Tab2: Luciferase constructs. Sequences and parameters of the proteins used in the split luciferase assay. Tab3: experimentally_validated_assemblies. List that specifies components of all linear assemblies shown in this work. Tab4: all_theoretical_assemblies. List of potential linear oligomers that could be assembled from the components shown in this work.

#### Data S2. S2_pdbs_and_scripts.tar.gz

Archive containing pdbs of components and assemblies shown in this work, as well as computational design scripts used to generate the heterodimers presented in this work.

---

**Figure s1.**
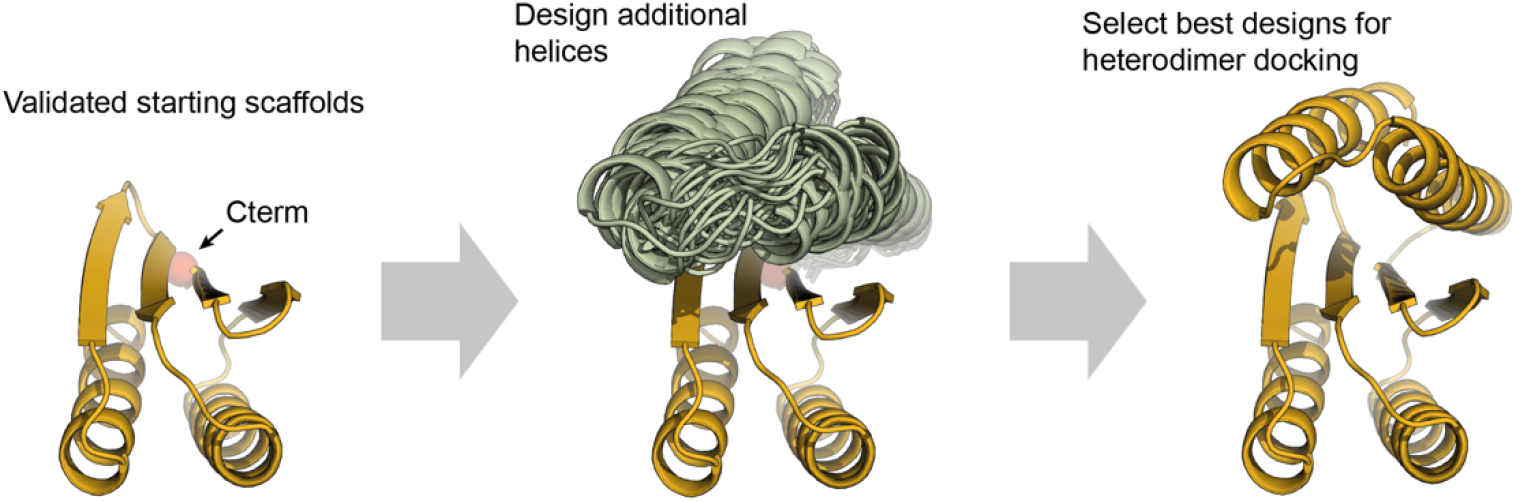
Modification Fold-it scaffolds. Fold-it scaffold 2003333_0006 (left) was expanded with 2 additional helices (middle) on its C-terminus via blueprint-based backbone generation. After backbone generation, the scaffold sequence was designed and the best scaffolds were selected (right) based on per residue rosetta energy and core packing.

**Figure s2.**
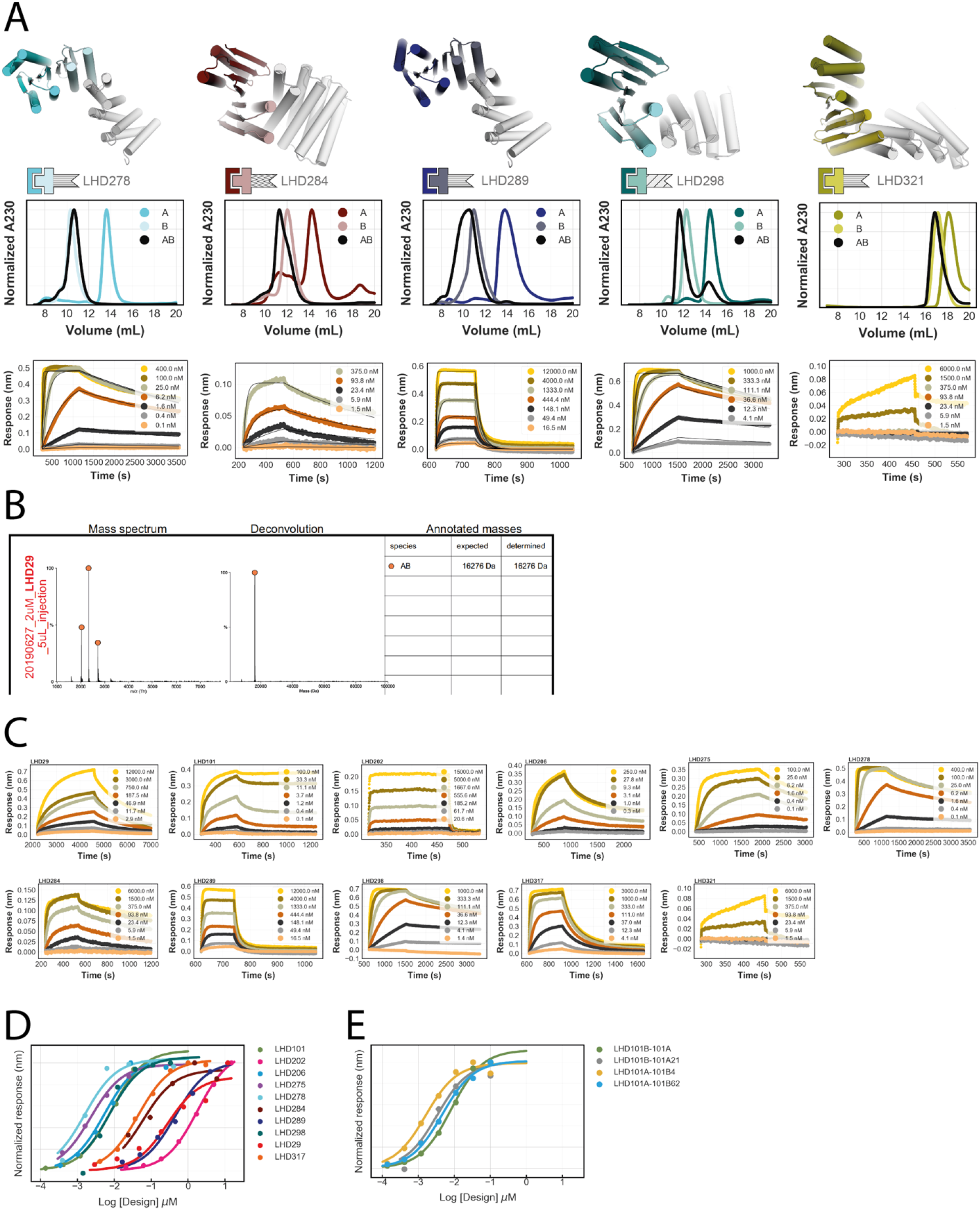
Characterization LHD binding in vitro. **A:** Designed models heterodimers (top row). Middle row, SEC binding experiments performed on a superdex 75 column. Bottom row, biolayer interferometry kinetic binding traces. **B:** Convoluted and deconvoluted native mass spectrums of the LHD29 heterodimer. **C:** Kinetic binding traces from BLI. Equilibrium responses were used to fit equilibrium binding curves **D:** Equilibrium binding curves of LHDs from biolayer interferometry binding assays with data from C. **E:** Equilibrium binding curves of unfused LHD101 protomers binding to rigid DHR fusions of LHD101B (DHR4 and 62) and LHD101A (DHR21). Biotinylated unfused protomers were immobilized on streptavidin coated biosensors.

**Figure s3.**
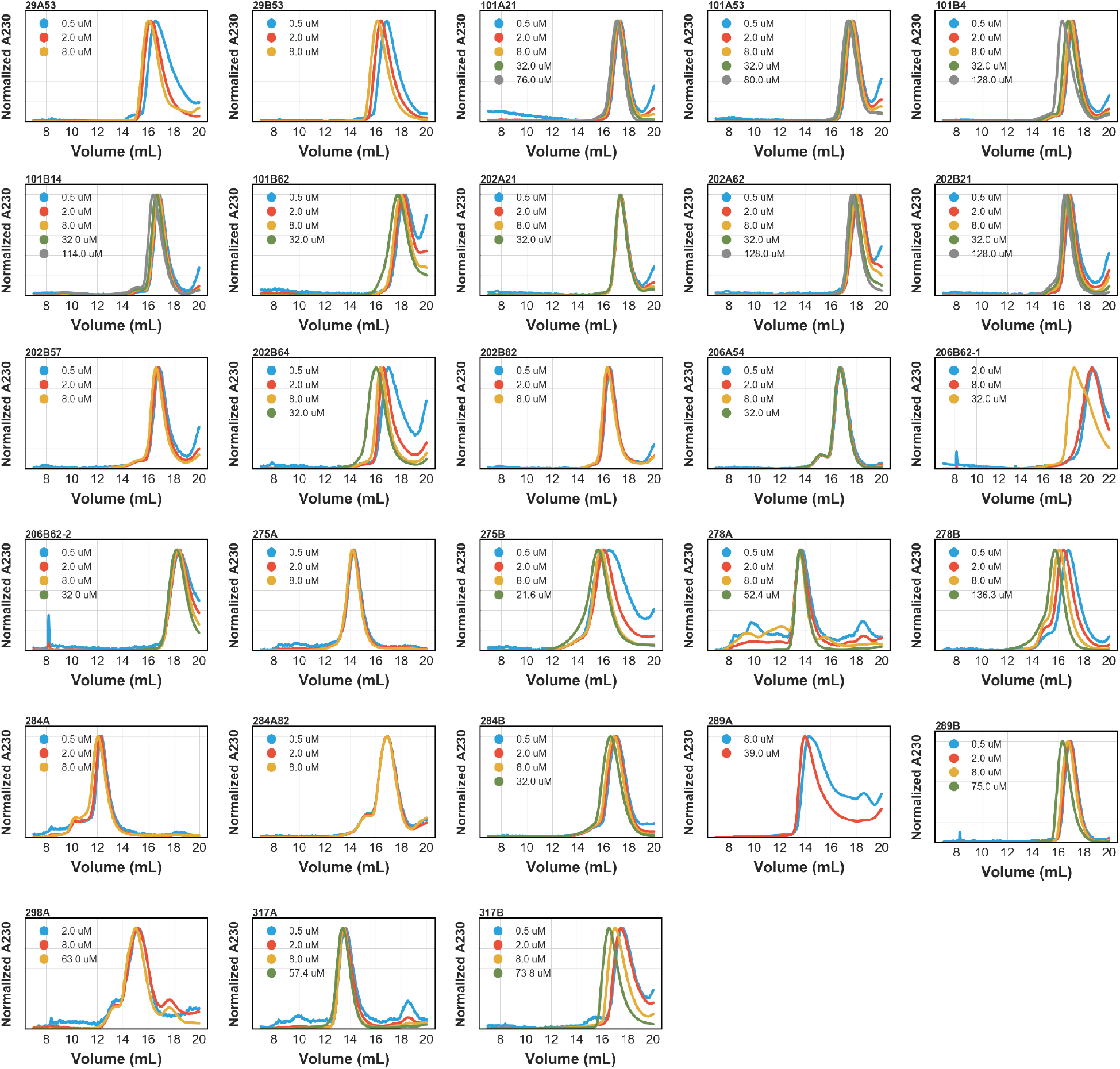
Oligomeric state of LHD protomers. SEC chromatograms of various LHD protomers titrated at indicated injection concentrations. All experiments were performed on a superdex 200 column except for LHDs 275A, 278A, 284A, 289A, 298A and 317A. These were run on a superdex 75 column.

**Figure s4.**
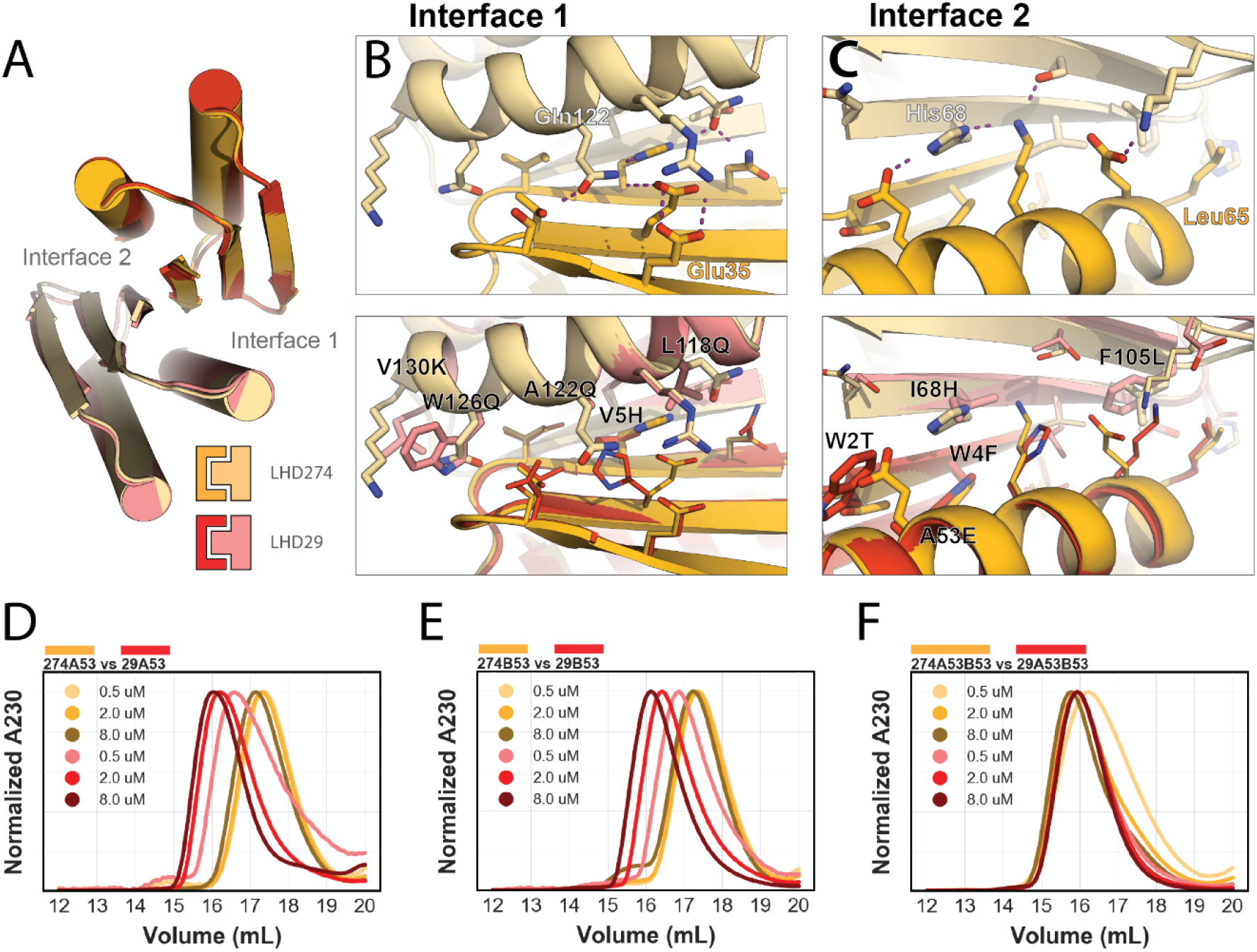
Redesign of LHD29. **A:** Superposition of a redesigned version of LHD29 designated LHD274 (yellows) and LHD29 (reds). Top, atomic view of interface 1 (**B)** region of LHD29 and interface 2 region (**C)**. Bottom panels, Overlay view of LHD29 and LHD274 at the corresponding region. Thick sticks indicate hydrophobic to polar substitutions. **D:** SEC superdex 200 titration of LHD29A and LHD274A fused to DHR53 at indicated concentrations. Fusion proteins were chosen for this assay for their enhanced absorbance at 230 nm compared to the much smaller unfused versions. **E:** SEC superdex 200 titration of LHD29B and LHD274B fused to DHR53 at indicated concentrations. **F:** Titration of the 29 and 274 complexes.

**Figure s5.**
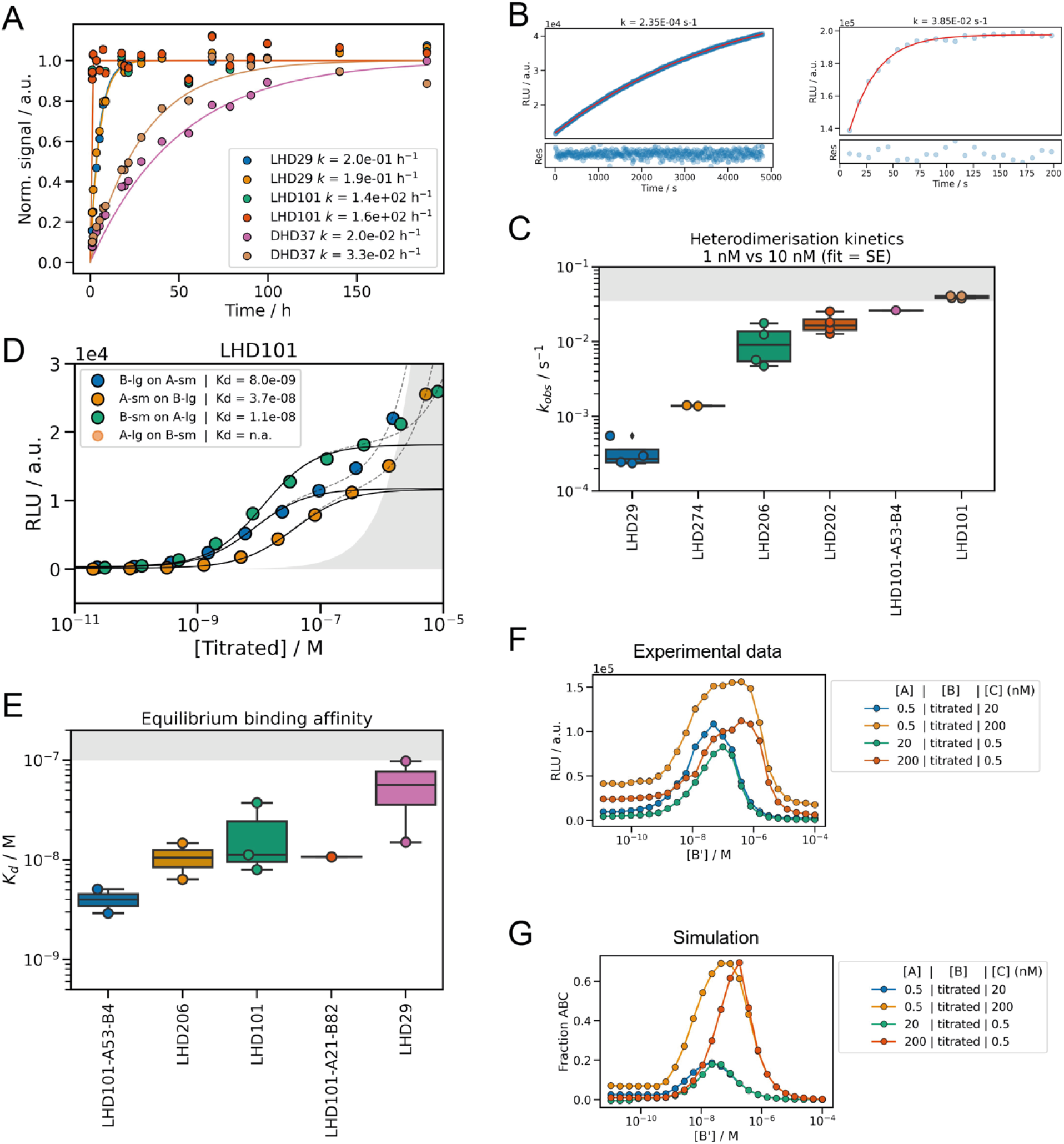
Characterization of binding interactions with a split luciferase reporter assay. Protein interactions were characterized by monitoring the reconstitution of split luciferase activity (smBiT:lgBiT) upon binding in buffer (from purified components; A, G-H) or lysate (B-F). **A** Comparison between the observed association kinetics of LHDs and designed helical hairpins (DHD37, previous work) under pseudo first-order conditions (1 nM *vs.* 10 nM). Reactions were monitored by taking manual time-points over the course of a week. The data was fitted to a single exponential decay function (solid line; rates are reported in the figure legend). **B** Example kinetic traces for the association of LHD29 (left) and LHD101 (right) in lysate. The data is shown in blue, and the single-exponential fits in red. Residuals to the fits are shown under each plot, and the rates are reported on top of each plot. **C** Summary statistics for association reactions performed under pseudo first-order conditions (1 nM *vs.* 10 nM) in lysate. Values are reported in Table S1. The grey shaded area indicates the limit of detection of the assay. **D** Example of equilibrium binding data collected in lysate (shown here for LHD101). Dashed lines are fits to the data, which includes a correction term to account for the intrinsic affinity of the split luciferase components (approximated by the grey shaded area). The binding curves (excluding the correction) are shown as solid black lines. The fitted *K*_d_ values are indicated in the figure legend. **E** Summary statistics for the equilibrium binding experiments performed in lysate. Values are reported in Table S2. **F,G** Equilibrium binding data (F) and simulation (G) for the ternary complex ABC. The data closely matches the prediction obtained from simulating the system with the affinities of each interface as measured in isolation (*K*_d_(LHD101) = 5 nM, *K*_d_(LHD29) = 50 nM), highlighting the modularity and transferability of LHD heterodimers.

**Figure s6.**
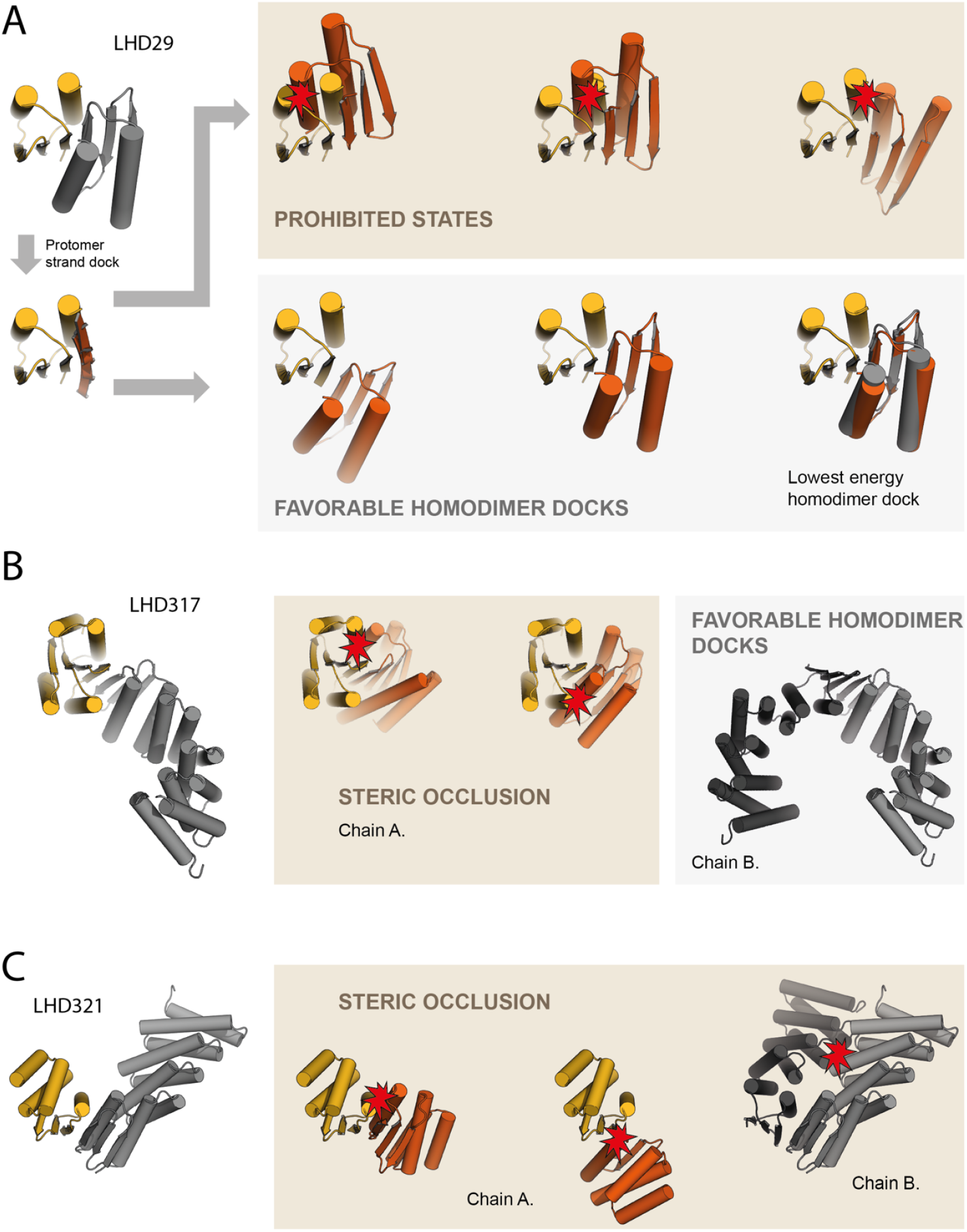
Homodimer docking. **A:** Example of homodimer docking. Homodimeric interaction most likely will occur on the edgestrand that forms the heterodimer. Strands are docked (orange) to the interface edgestrand of a protomer (yellow) of a given heterodimer. Another copy of the same protomer (orange) is then aligned along the docked edgestrand to create a homodimeric docked complex. Most complexes clash indicating homodimerization is unfavorable (top row). Some docks do not clash (bottom row) but have limited interaction surface area making homodimerization unlikely. In some cases homodimer docks i.e. LHD29 have similar interactions energies as the heterodimer (bottom right). These docks are likely to form homodimers. **B:** Homodimer docking of LHD317 protomers shows that secondary structure elements prevent LHD317A homodimerization via steric occlusion whereas 317B homodimers are more favorable. **C:** Designed secondary structure elements in both protomors of LHD321 prevent homodimerization

**Figure s7.**
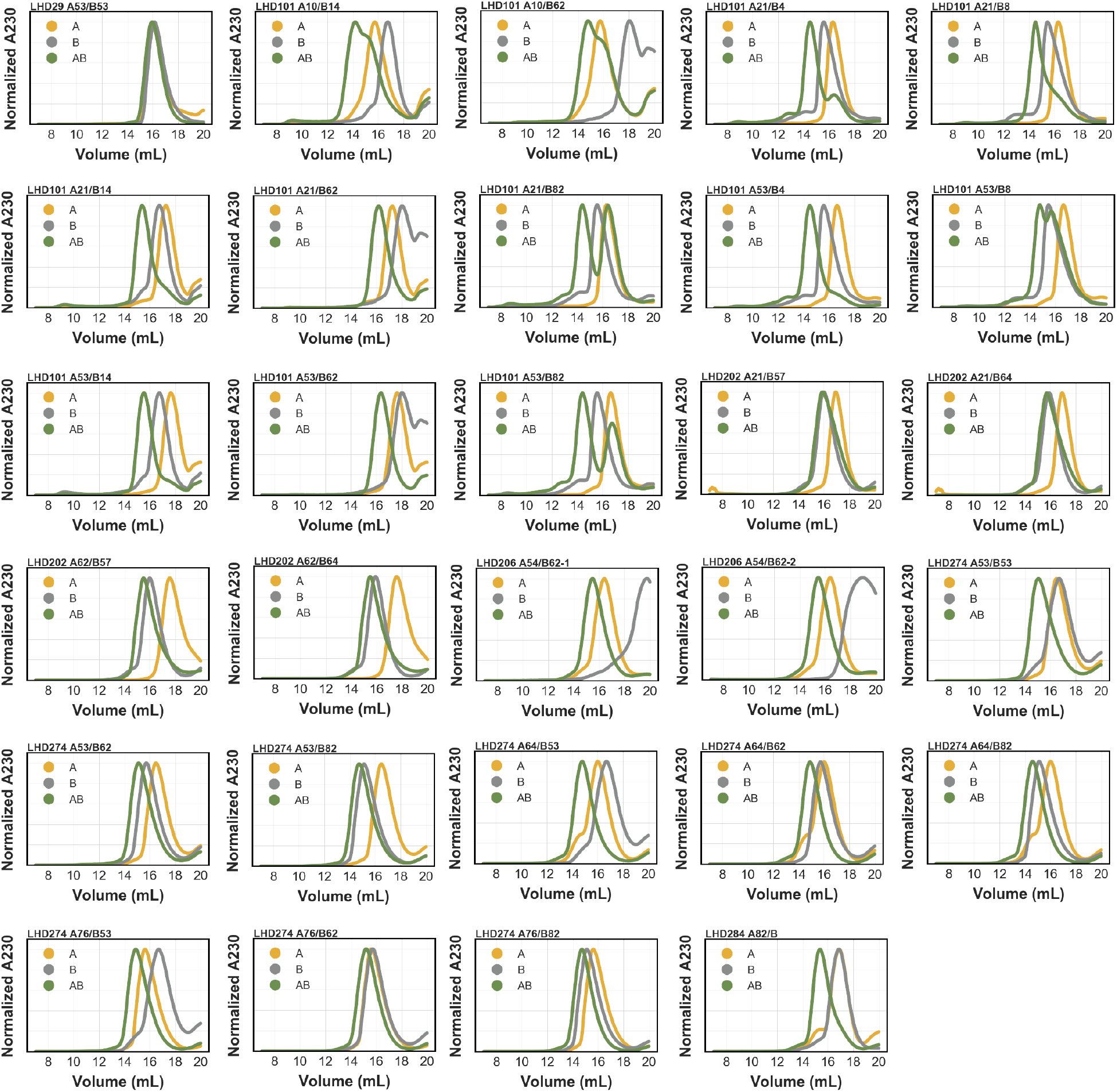
LHD fusion binding assays. Superdex 200 binding assays of LHD fusion proteins.

**Figure s8.**
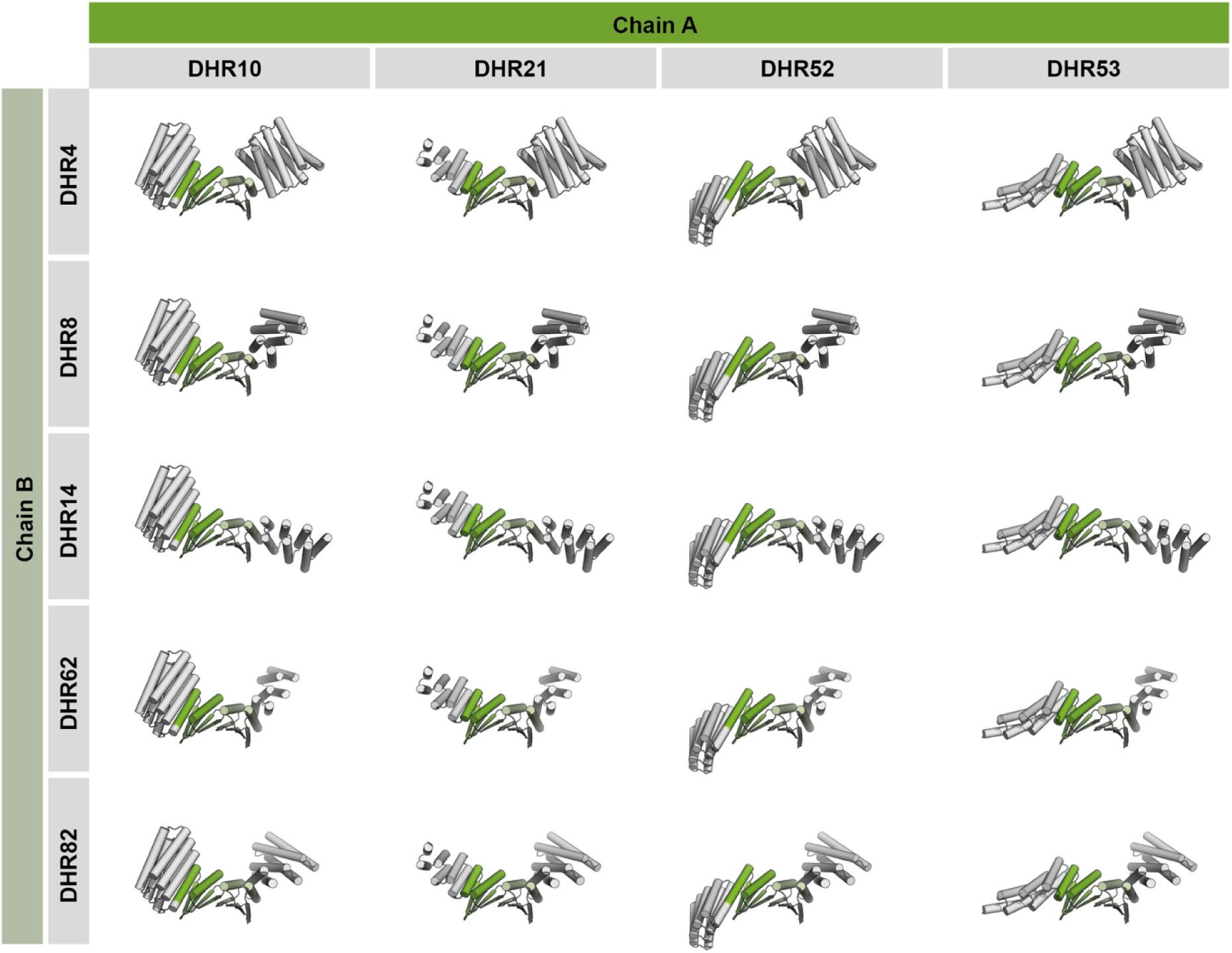
Models LHD101 fusion complexes. Designed models of all possible 20 complexes involving LHD101 fusions. DHRs colored in white and LHD base core protomers in greens. Combinations with unfused protomers (10 complexes) are not shown.

**Figure s9.**
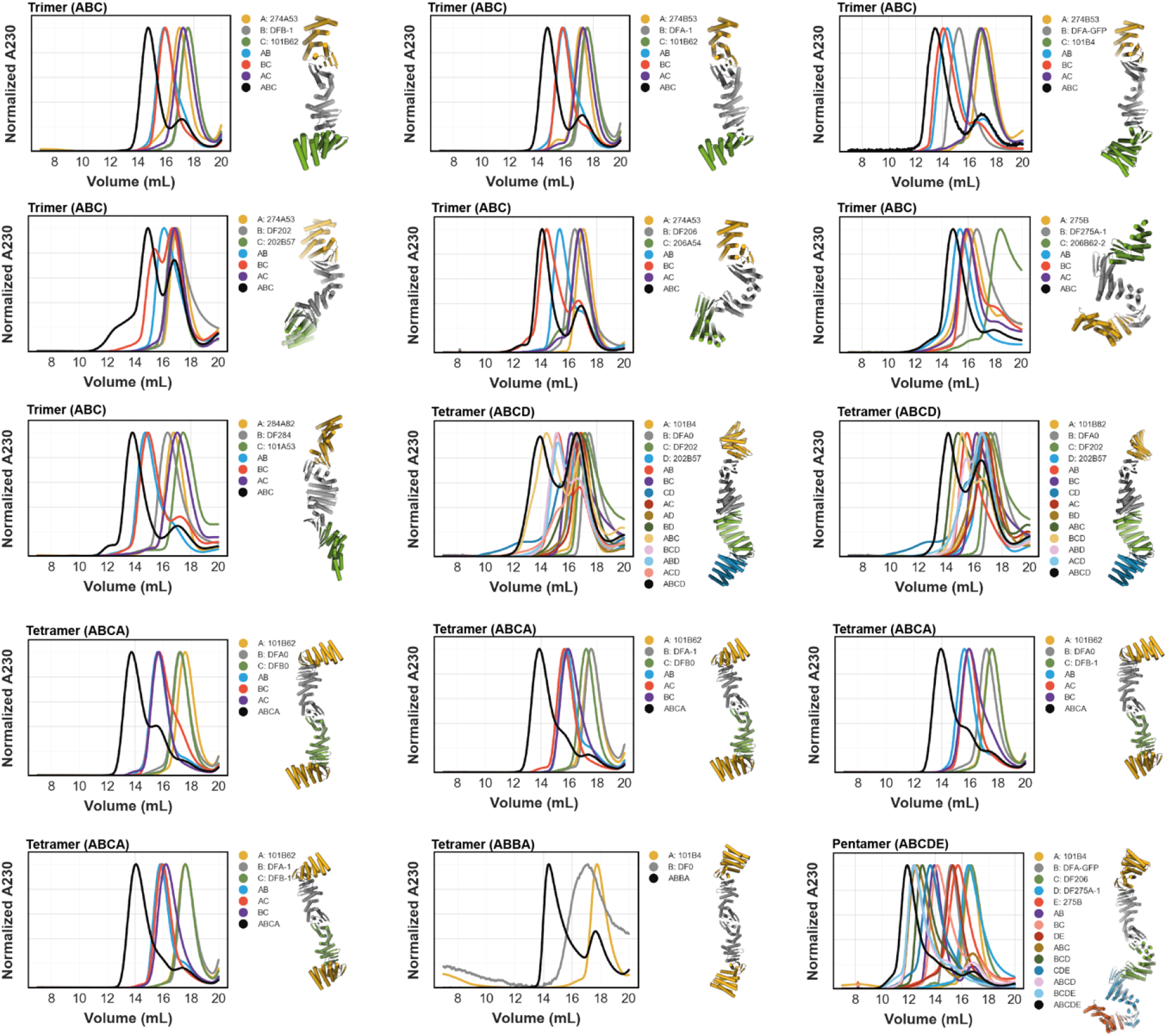
SEC binding assays linear hetero-oligomers. Superdex 200 chromatograms of various linear assemblies and their control sub-assemblies. Designed models of the target assembly (black chromatogram) are shown right of the graphs

**Figure s10.**
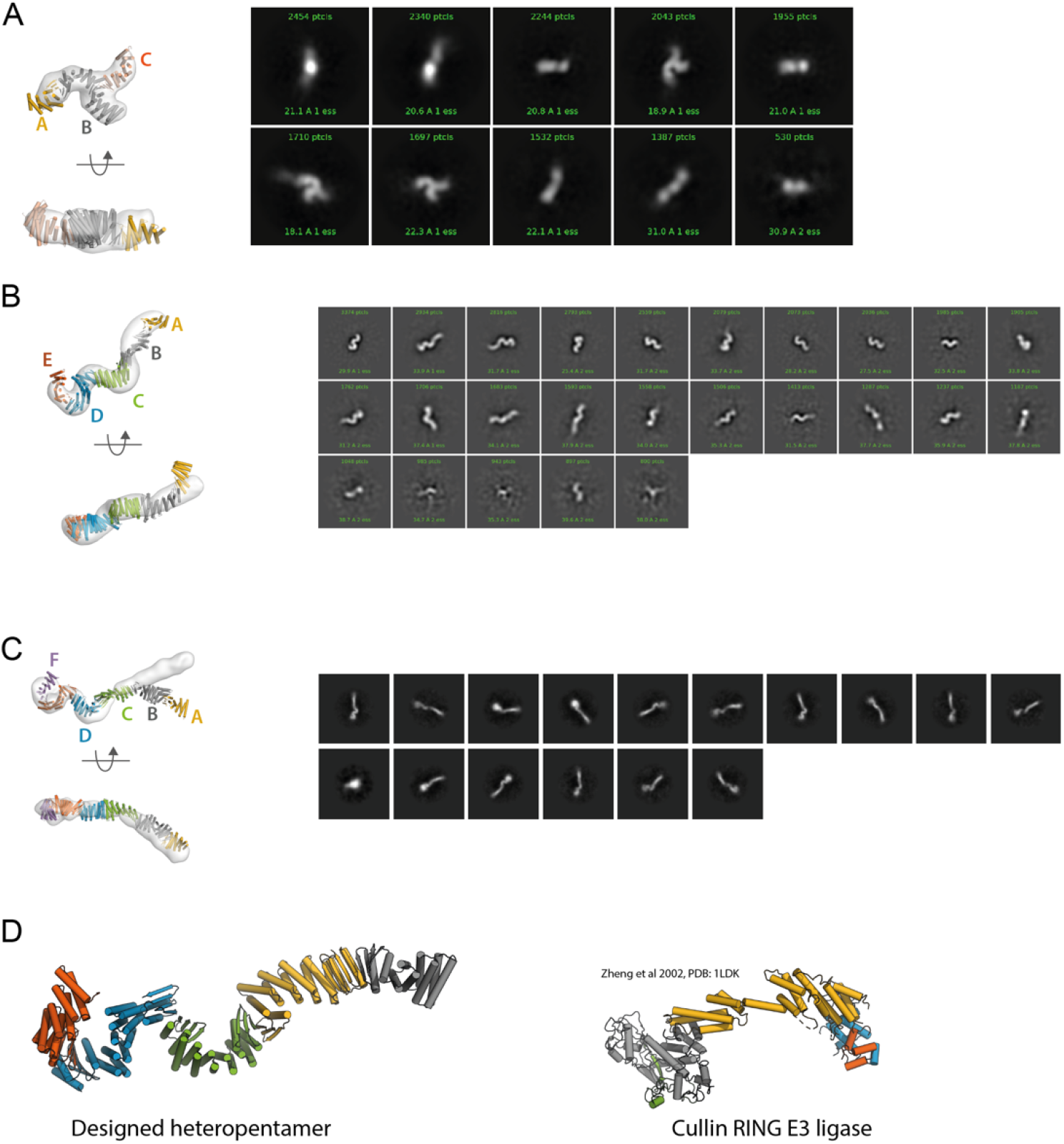
Negative stain EM class averages and 3D reconstructions hetero-oligomers. **A:** Heterotrimer (ABC) consisting of LHD274A53 (A), linear connector DFx (B) and LHD317B (C). **B:** Heteropentamer (ABCDE) consisting of 101B4 (A), DFA0 (B), DF206 (C), DF275A-1 (D) and 275B (E). **C:** Heterohexamer consisting of 284A82 (A), DF284B (B), DFA0 (C), DF206 (D), DF275A-1 (E) and 275B (F). **D:** Comparison between designed heteropentamer (left) and the Cul1-Rbx1-Skp1-F box^Skp2^ SCF ubiquitin ligase complex (right) <u>(Zheng et al. 2002).</u>

**Figure s11.**
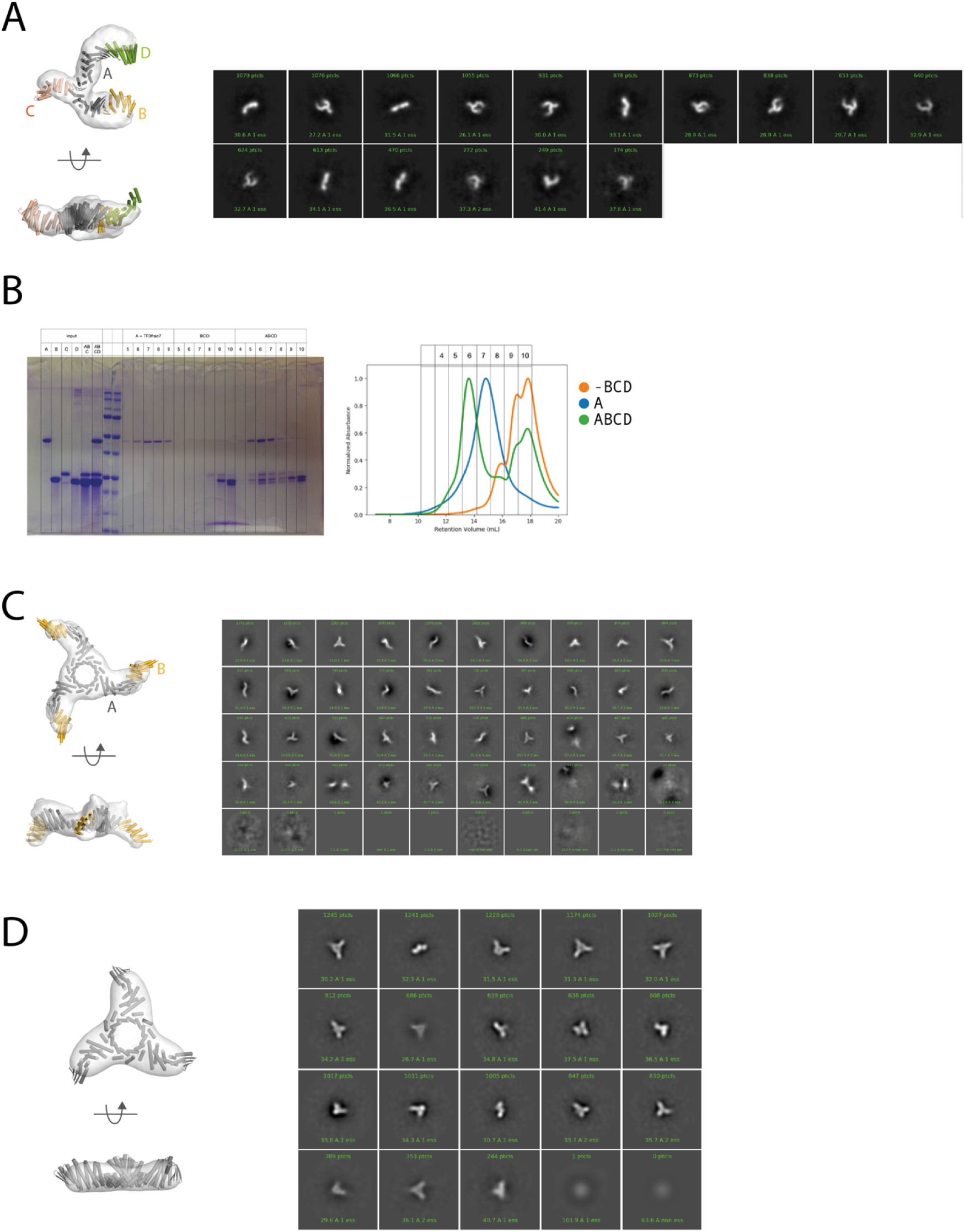
Non-linearly arranged assemblies. **A:** Class averages and 3D reconstruction of a branched tetramer (ABCD) consisting of trivalent connector TF10 (A), LHD274A53 (B), LHD317B (C) and LHD101B62 (D). **B:** SEC and corresponding SDS-PAGE analysis of a branched tetramer consisting of trivalent connector TF3 (A), LHD274A53 (B), LHD275B (C) and LHD101B62. **C and D:** Class averages and 3D reconstruction of the C3-Hub bound to LHD101A53 and by itself.

**Figure s12.**
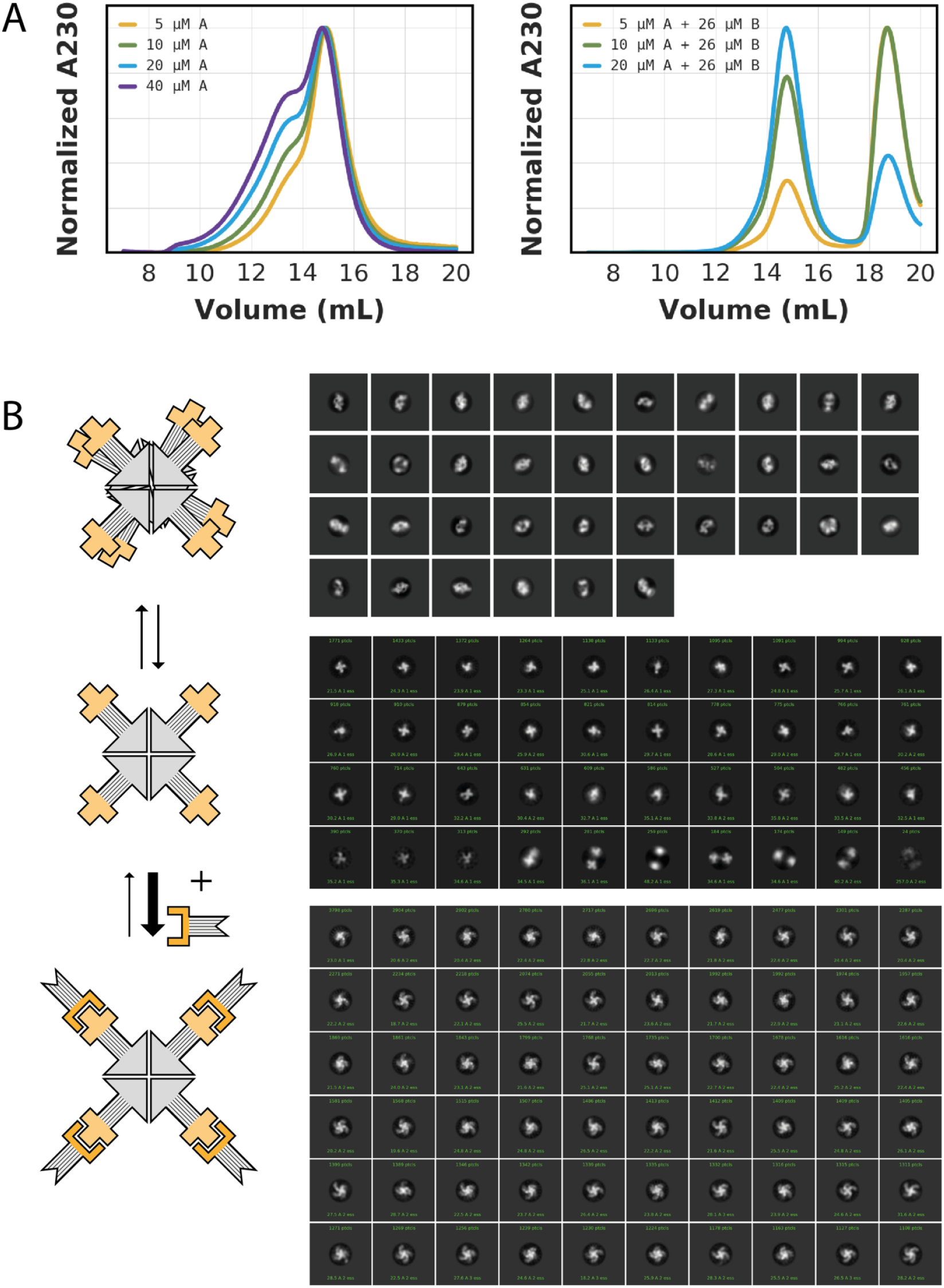
Characterization of C4 hetero-oligomers. **A:** SEC traces of the C4-symmetric hub at different concentrations without binding partner (left) and with a constant concentration of binding partner (right). Concentrations are given per monomer (5 µM corresponds to 1.25 µM tetramer). **B:** Schematic representations (left; (dark grey: C4 hub, gold: binding partner) and negative stain EM class averages (right) of the C4-symmetric hub without (top, center) and with (bottom) binding partner. In absence of the binding partner, the C4 hub exists in equilibrium between a higher order complex (top) and the designed C4 complex (center).

**Figure s13.**
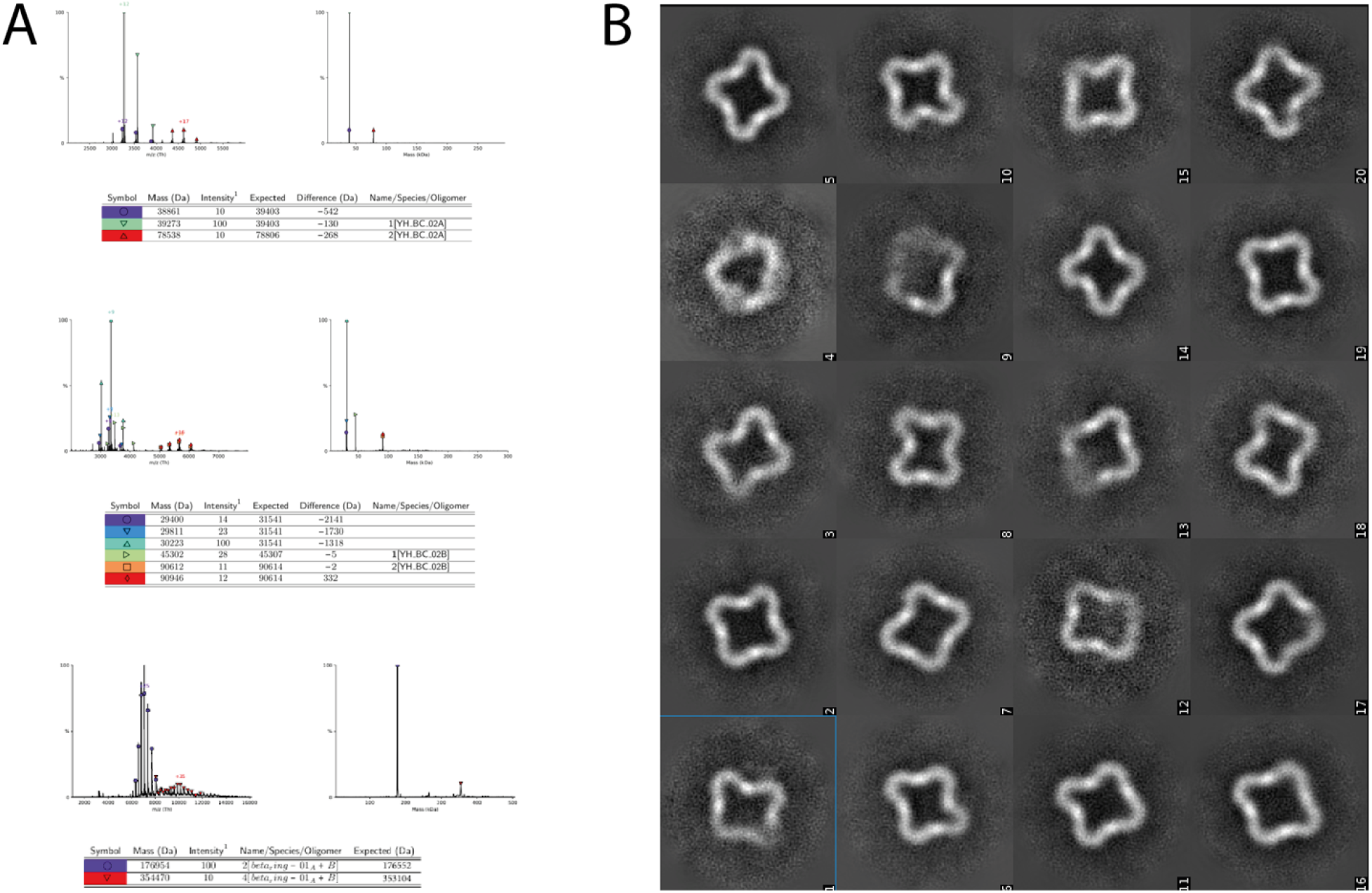
Characterization of the closed C4-symmetric ring. **A:** Convoluted and deconvoluted native mass spectrums of the two component C4-symmetrical ring and constituent components. **B:** Negative stain EM class averages of the closed C4-symmetric ring shown in Fig. 4D

**Figure s14.**
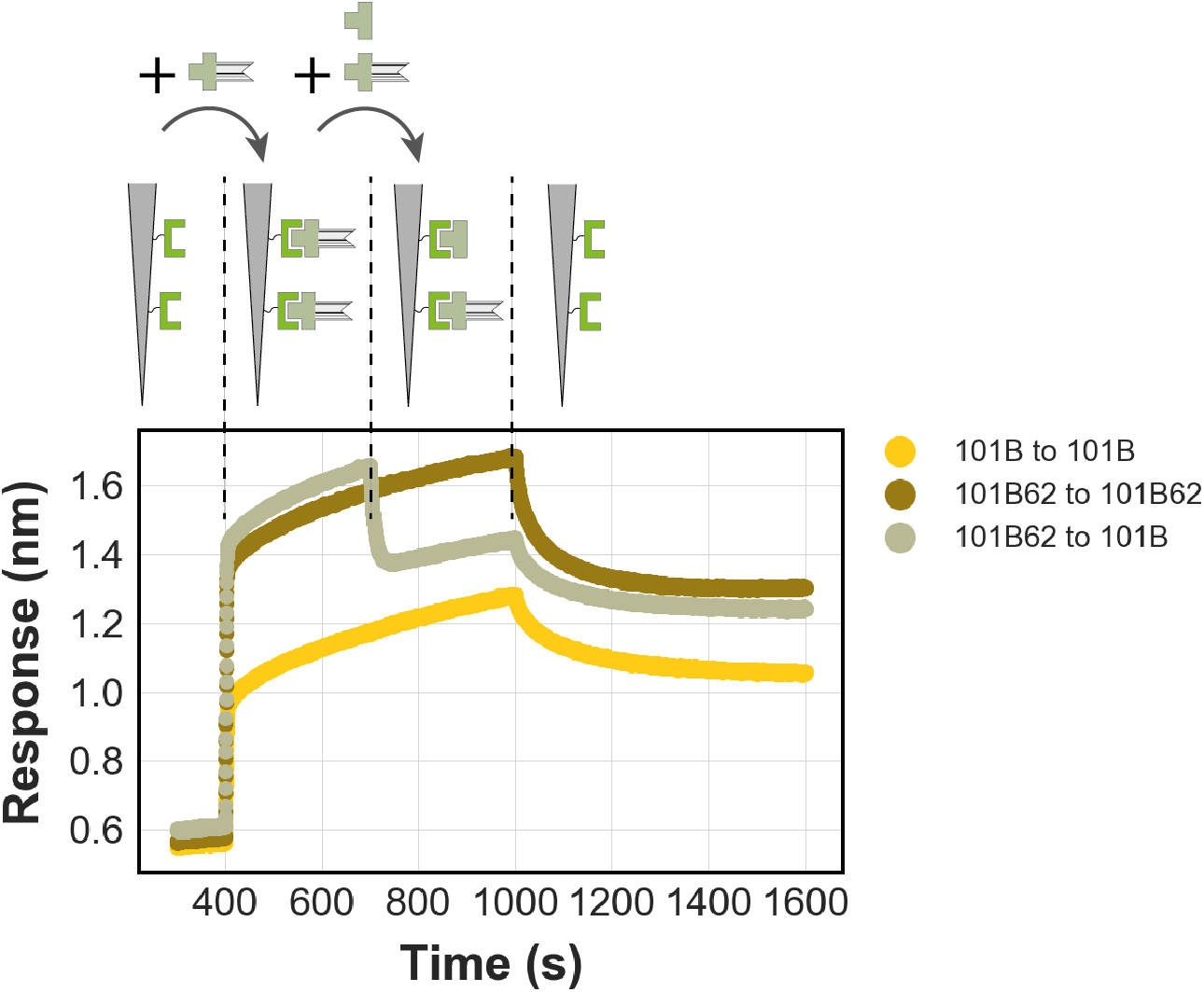
Biolayer interferometry subunit exchange. Biotinylated LHD101 that is immobilized to streptavidin biosensors binds rigid fusion variant LHD101B62. Biosensors were next dipped into a solution containing equimolar amounts of LHD101B62 and unfused 101B at saturating concentrations. The binding response of this reaction is in between controls (brown and yellow) indicating subunit exchange takes place.

**Table S1.** Fitted values biolayer interferometry binding assays

| Design | Steady state fits |  | Kinetic fits |  |  |  |  |
| --- | --- | --- | --- | --- | --- | --- | --- |
| | $K_D$ (nM) | R-sqr | $K_D$ (nM) | $k_{on}$ ($M^{-1} s^{-1}$) | $k_{off}$ ($s^{-1}$) | chi-sqr | R-sqr |
| **LHD29<sup>1</sup>** | $310 \pm 120$ | 0.91 | $985 \pm 6.0$ | $6.9 \cdot 10^2 \pm 4$ | $6.8 \cdot 10^{-4} \pm 1.1 \cdot 10^{-6}$ | 6.7 | 0.98 |
| **LHD101** | $9.5 \pm 0.76$ | 0.99 | $1.9 \pm 0.04$ | $2.2 \cdot 10^6 \pm 4.0 \cdot 10^4$ | $4.3 \cdot 10^{-3} \pm 2.1 \cdot 10^{-5}$ | 0.21 | 0.97 |
| **LHD202** | $2400 \pm 170$ | 0.99 | $4800 \pm 250$ | $6.0 \cdot 10^4 \pm 3.0 \cdot 10^3$ | $2.9 \cdot 10^{-1} \pm 0.05$ | 0.03 | 0.99 |
| **LHD206** | $8.4 \pm 1.6$ | 0.97 | $2.8 \pm 0.02$ | $2.7 \cdot 10^5 \pm 1.9 \cdot 10^3$ | $7.5 \cdot 10^{-4} \pm 2.2 \cdot 10^{-6}$ | 0.8 | 0.99 |
| **LHD274** | nd | nd | nd | nd | nd | nd | nd |
| **LHD275** | $4.5 \pm 0.22$ | 0.99 | $2.9 \pm 0.01$ | $1.4 \cdot 10^5 \pm 4.6 \cdot 10^2$ | $4.1 \cdot 10^{-4} \pm 1.1 \cdot 10^{-6}$ | 0.76 | 0.99 |
| **LHD278** | $3.4 \pm 0.69$ | 0.98 | $0.8 \pm 0.003$ | $2.9 \cdot 10^5 \pm 1 \cdot 10^3$ | $2.2 \cdot 10^{-4} \pm 3.6 \cdot 10^{-7}$ | 2.8 | 0.99 |
| **LHD284** | $97 \pm 13$ | 0.99 | $8.9 \pm 0.13$ | $1.3 \cdot 10^5 \pm 1.7 \cdot 10^3$ | $1.2 \cdot 10^{-3} \pm 6.7 \cdot 10^{-6}$ | 0.06 | 0.99 |
| **LHD289** | $610 \pm 120$ | 0.97 | $1080 \pm 39$ | $5.3 \cdot 10^4 \pm 1.9 \cdot 10^3$ | $5.7 \cdot 10^{-2} \pm 5.8 \cdot 10^{-4}$ | 0.99 | 0.99 |
| **LHD298** | $16 \pm 3$ | 0.97 | $3.5 \pm 0.01$ | $6.4 \cdot 10^4 \pm 1.0 \cdot 10^2$ | $2.2 \cdot 10^{-4} \pm 5.9 \cdot 10^{-7}$ | 6.4 | 0.99 |
| **LHD317** | $56 \pm 2.3$ | 0.99 | $34.7 \pm 0.05$ | $1.5 \cdot 10^5 \pm 2.1 \cdot 10^3$ | $5.1 \cdot 10^{-3} \pm 1.6 \cdot 10^{-5}$ | 4.7 | 0.99 |
| **LHD321** | nd | nd | nd | nd | nd | nd | nd |

<sup>1</sup>Homodimerization of both LHD29 protomers under BLI conditions make $K_D$ determination unreliable. $K_D$ from split luciferase assay (Fig. S5) is more reliable as the experiment was performed under dilute conditions where homodimerization is minimized. nd: not determined

**Table S2.** Fitted rate constants for heterodimerization reactions performed at 1 nM *vs.* 10 nM in lysate. Errors indicate standard deviations.

| **Design** | **$k_{\text{obs}}$ (s<sup>-1</sup>)** |
| --- | --- |
| DHD37*,<sup>1</sup> | $7 \pm 3 \cdot 10^{-6}$ |
| LHD29 | $3 \pm 1 \cdot 10^{-4}$ |
| LHD29* | $5.5 \pm 2 \cdot 10^{-5}$ |
| LHD274 | $1.40 \pm 0.01 \cdot 10^{-3}$ |
| LHD206 | $1.0 \pm 0.5 \cdot 10^{-2}$ |
| LHD202 | $1.8 \pm 0.5 \cdot 10^{-2}$ |
| LHD101-A53-B4 | $2.6 \cdot 10^{-2}$ |
| LHD101 | $4.0 \pm 0.1 \cdot 10^{-2}$ |
| LHD101* | $4.2 \pm 0.4 \cdot 10^{-2}$ |

<sup>1</sup>(Chen et al. 2019). \* Experiments performed with purified proteins, and reactions monitored by taking manual time-points as described in Materials and Methods and Supplementary Materials and Methods .

**Table S3.** Fitted equilibrium dissociation constants for binding curves collected in lysate. Errors indicate standard deviations.

| **Design** | **$K_d$ (M)** |
| --- | --- |
| LHD101 | $2 \pm 1 \cdot 10^{-8}$ |
| LHD206 | $1.1 \pm 0.4 \cdot 10^{-8}$ |
| LHD101-A21-B82 | $1.1 \cdot 10^{-8}$ |
| LHD29 | $6 \pm 4 \cdot 10^{-8}$ |
| LHD101-A53-B4 | $4 \pm 1 \cdot 10^{-9}$ |

**Table S4.** Crystallographic data collection and refinement.

|  | **LHD29 (PDB:6WMK)** | **LHD29A53/B53 (PDB: 7MWQ)** | **LHD101A53/B4 (PDB: 7MWR)** |
| --- | --- | --- | --- |
| **Data Collection** |  |  |  |
| Space group | $P 2_1$ | $P1$ | $P 2_1 2_1 2_1$ |
| Cell dimensions |  |  |  |
| $a, b, c$ (Å) | 56.07, 38.17, 60.37 | 61.31, 73.45, 4.14 | 45.40, 99.77, 122.09 |
| $\alpha, \beta, \gamma$ (°) | 90, 98.26, 90 | 108.39, 106.70, 110.15 | 90.0, 90.0, 90.0 |
| Resolution (Å) | 38.03 – 2.20 (2.42 - 2.20) | 51.56 - 2.56 (2.65 - 2.56) | 42.56 - 2.2 (2.27 - 2.20) |
| $R_{merge}$ (%) | 7 (56.9) | 8.3 (82.8) | 3.1 (49.2) |
| $R_{pim}$ (%) | 4.6 (36.5) | 6.6 (69.5) | 3.1 (49.2) |
| $I/\sigma(I)$ | 6.3 (1.4) | 4.7 (1.07) | 15.9 (1.6) |
| $CC_{1/2}$ | 0.995 (0.705) | 0.991(0.651) | 0.999 (0.757) |
| Completeness (%) | 94.2 (99.2) | 97.9 (93.4) | 99.8 (99.0) |
| Redundancy | 3.3 (3.3) | 2.3 (2.4) | 2.0 (2.0) |
| **Refinement** |  |  |  |
| Resolution (Å) | 38.03 – 2.20 (2.42 - 2.20) | 51.56 - 2.56 (2.65 - 2.56) | 42.56 - 2.2 (2.27 - 2.20) |
| No. reflections | 12330 | 32540 | 28939 |
| $R_{work} / R_{free}$ (%) | 25.3 / 28.3 (29.9 / 37.1) | 23.2 / 26.9 (36.9 / 41.9) | 21.1 /25.2 (40.6 /40.1) |
| No. atoms | 2154 | 6384 | 3514 |
| Protein | 2105 | 6370 | 11544 |
| Ligand | n/a | n/a | 7 |
| Water | 49 | 14 | 82 |
| Ramachandran Favored/allowed Outlier (%) | 96.80/3.20 | 98.64 / 1.11<br>0.25 | 97.77 /2.23<br>0.00 |
| R.m.s. deviations |  |  |  |
| Bond lengths (Å) | 0.002 | 0.002 | 0.002 |
| Bond angles (°) | 0.394 | 0.40 | 0.41 |
| $B_{factors}$ (Å<sup>2</sup>) | | | |
| Protein | 55.00 | 75.64 | 52.36 |
| Ligand | n/a | n/a | 78.04 |
| Water | 42.13 | 53.18 | 53.31 |

## References and Notes

1. S. E. Tusk, N. J. Delalez, R. M. Berry, Subunit Exchange in Protein Complexes. J. Mol. Biol. 430, 4557–4579 (2018).

2. C. Engel, S. Neyer, P. Cramer, Distinct Mechanisms of Transcription Initiation by RNA Polymerases I and II. Annu. Rev. Biophys. 47, 425–446 (2018).

3. P. M. J. Burgers, T. A. Kunkel, Eukaryotic DNA Replication Fork. Annu. Rev. Biochem. 86, 417–438 (2017).

4. S. Gonen, F. DiMaio, T. Gonen, D. Baker, Design of ordered two-dimensional arrays mediated by noncovalent protein-protein interfaces. Science. 348, 1365–1368 (2015).

5. Y. Hsia, J. B. Bale, S. Gonen, D. Shi, W. Sheffler, K. K. Fong, U. Nattermann, C. Xu, P.-S. Huang, R. Ravichandran, S. Yi, T. N. Davis, T. Gonen, N. P. King, D. Baker, Design of a hyperstable 60-subunit protein dodecahedron. [corrected]. Nature. 535, 136–139 (2016).

6. N. P. King, J. B. Bale, W. Sheffler, D. E. McNamara, S. Gonen, T. Gonen, T. O. Yeates, D. Baker, Accurate design of co-assembling multi-component protein nanomaterials. Nature. 510, 103–108 (2014).

7. Y. Hsia, R. Mout, W. Sheffler, N. I. Edman, I. Vulovic, Y.-J. Park, R. L. Redler, M. J. Bick, A. K. Bera, A. Courbet, A. Kang, T. J. Brunette, U. Nattermann, E. Tsai, A. Saleem, C. M. Chow, D. Ekiert, G. Bhabha, D. Veesler, D. Baker, Design of multi-scale protein complexes by hierarchical building block fusion. Nat. Commun. 12, 2294 (2021).

8. A. J. Ben-Sasson, J. L. Watson, W. Sheffler, M. C. Johnson, A. Bittleston, L. Somasundaram, J. Decarreau, F. Jiao, J. Chen, I. Mela, A. A. Drabek, S. M. Jarrett, S. C. Blacklow, C. F. Kaminski, G. L. Hura, J. J. De Yoreo, J. M. Kollman, H. Ruohola-Baker, E. Derivery, D. Baker, Design of biologically active binary protein 2D materials. Nature. 589, 468–473 (2021).

9. R. Divine, H. V. Dang, G. Ueda, J. A. Fallas, I. Vulovic, W. Sheffler, S. Saini, Y. T. Zhao, I. X. Raj, P. A. Morawski, M. F. Jennewein, L. J. Homad, Y.-H. Wan, M. R. Tooley, F. Seeger, A. Etemadi, M. L. Fahning, J. Lazarovits, A. Roederer, A. C. Walls, L. Stewart, M. Mazloomi, N. P. King, D. J. Campbell, A. T. McGuire, L. Stamatatos, H. Ruohola-Baker, J. Mathieu, D. Veesler, D. Baker, Designed proteins assemble antibodies into modular nanocages. Science. 372 (2021), doi:10.1126/science.abd9994.

10. Z. Chen, S. E. Boyken, M. Jia, F. Busch, D. Flores-Solis, M. J. Bick, P. Lu, Z. L. VanAernum, A. Sahasrabuddhe, R. A. Langan, S. Bermeo, T. J. Brunette, V. K. Mulligan, L. P. Carter, F. DiMaio, N. G. Sgourakis, V. H. Wysocki, D. Baker, Programmable design of orthogonal protein heterodimers. Nature. 565, 106–111 (2019).

11. S. E. Boyken, Z. Chen, B. Groves, R. A. Langan, G. Oberdorfer, A. Ford, J. M. Gilmore, C. Xu, F. DiMaio, J. H. Pereira, B. Sankaran, G. Seelig, P. H. Zwart, D. Baker, De novo design of protein homo-oligomers with modular hydrogen-bond network-mediated specificity. Science. 352, 680–687 (2016).

12. Z. Chen, R. D. Kibler, A. Hunt, F. Busch, J. Pearl, M. Jia, Z. L. VanAernum, B. I. M. Wicky, G. Dods, H. Liao, M. S. Wilken, C. Ciarlo, S. Green, H. El-Samad, J. Stamatoyannopoulos, V. H. Wysocki, M. C. Jewett, S. E. Boyken, D. Baker, De novo design of protein logic gates. Science. 368, 78–84 (2020).

13. H. Gradišar, R. Jerala, De novo design of orthogonal peptide pairs forming parallel coiled-coil heterodimers. J. Pept. Sci. 17, 100–106 (2011).

14. C. L. Edgell, A. J. Smith, J. L. Beesley, N. J. Savery, D. N. Woolfson, De Novo Designed Protein-Interaction Modules for In-Cell Applications. ACS Synth. Biol. 9, 427–436 (2020).

15. A. Leaver-Fay, R. Jacak, P. B. Stranges, B. Kuhlman, A generic program for multistate protein design. PLoS One. 6, e20937 (2011).

16. A. Leaver-Fay, K. J. Froning, S. Atwell, H. Aldaz, A. Pustilnik, F. Lu, F. Huang, R. Yuan, S. Hassanali, A. K. Chamberlain, J. R. Fitchett, S. J. Demarest, B. Kuhlman, Computationally Designed Bispecific Antibodies using Negative State Repertoires. Structure. 24, 641–651 (2016).

17. S. J. Fleishman, D. Baker, Role of the biomolecular energy gap in protein design, structure, and evolution. Cell. 149, 262–273 (2012).

18. D. D. Sahtoe, A. Coscia, N. Mustafaoglu, L. M. Miller, D. Olal, I. Vulovic, T.-Y. Yu, I. Goreshnik, Y.-R. Lin, L. Clark, F. Busch, L. Stewart, V. H. Wysocki, D. E. Ingber, J. Abraham, D. Baker, Transferrin receptor targeting by de novo sheet extension. Proc. Natl. Acad. Sci. U. S. A. 118 (2021), doi:10.1073/pnas.2021569118.

19. P. B. Stranges, M. Machius, M. J. Miley, A. Tripathy, B. Kuhlman, Computational design of a symmetric homodimer using β-strand assembly. Proc. Natl. Acad. Sci. U. S. A. 108, 20562–20567 (2011).

20. H. Remaut, G. Waksman, Protein-protein interaction through beta-strand addition. Trends Biochem. Sci. 31, 436–444 (2006).

21. B. Koepnick, J. Flatten, T. Husain, A. Ford, D.-A. Silva, M. J. Bick, A. Bauer, G. Liu, Y. Ishida, A. Boykov, R. D. Estep, S. Kleinfelter, T. Nørgård-Solano, L. Wei, F. Players, G. T. Montelione, F. DiMaio, Z. Popović, F. Khatib, S. Cooper, D. Baker, De novo protein design by citizen scientists. Nature. 570, 390–394 (2019).

22. T. J. Brunette, M. J. Bick, J. M. Hansen, C. M. Chow, J. M. Kollman, D. Baker, Modular repeat protein sculpting using rigid helical junctions. Proc. Natl. Acad. Sci. U. S. A. 117, 8870–8875 (2020).

23. Y.-R. Lin, N. Koga, R. Tatsumi-Koga, G. Liu, A. F. Clouser, G. T. Montelione, D. Baker, Control over overall shape and size in de novo designed proteins. Proc. Natl. Acad. Sci. U. S. A. 112, E5478–85 (2015).

24. N. Koga, R. Tatsumi-Koga, G. Liu, R. Xiao, T. B. Acton, G. T. Montelione, D. Baker, Principles for designing ideal protein structures. Nature. 491, 222–227 (2012).

25. J. K. Leman, B. D. Weitzner, S. M. Lewis, J. Adolf-Bryfogle, N. Alam, R. F. Alford, M. Aprahamian, D. Baker, K. A. Barlow, P. Barth, B. Basanta, B. J. Bender, K. Blacklock, J. Bonet, S. E. Boyken, P. Bradley, C. Bystroff, P. Conway, S. Cooper, B. E. Correia, B. Coventry, R. Das, R. M. De Jong, F. DiMaio, L. Dsilva, R. Dunbrack, A. S. Ford, B. Frenz, D. Y. Fu, C. Geniesse, L. Goldschmidt, R. Gowthaman, J. J. Gray, D. Gront, S. Guffy, S. Horowitz, P.-S. Huang, T. Huber, T. M. Jacobs, J. R. Jeliazkov, D. K. Johnson, K. Kappel, J. Karanicolas, H. Khakzad, K. R. Khar, S. D. Khare, F. Khatib, A. Khramushin, I. C. King, R. Kleffner, B. Koepnick, T. Kortemme, G. Kuenze, B. Kuhlman, D. Kuroda, J. W. Labonte, J. K. Lai, G. Lapidoth, A. Leaver-Fay, S. Lindert, T. Linsky, N. London, J. H. Lubin, S. Lyskov, J. Maguire, L. Malmström, E. Marcos, O. Marcu, N. A. Marze, J. Meiler, R. Moretti, V. K. Mulligan, S. Nerli, C. Norn, S. Ó’Conchúir, N. Ollikainen, S. Ovchinnikov, M. S. Pacella, X. Pan, H. Park, R. E. Pavlovicz, M. Pethe, B. G. Pierce, K. B. Pilla, B. Raveh, P. D. Renfrew, S. S. R. Burman, A. Rubenstein, M. F. Sauer, A. Scheck, W. Schief, O. Schueler-Furman, Y. Sedan, A. M. Sevy, N. G. Sgourakis, L. Shi, J. B. Siegel, D.-A. Silva, S. Smith, Y. Song, A. Stein, M. Szegedy, F. D. Teets, S. B. Thyme, R. Y.-R. Wang, A. Watkins, L. Zimmerman, R. Bonneau, Macromolecular modeling and design in Rosetta: recent methods and frameworks. Nat. Methods. 17, 665–680 (2020).

26. B. Coventry, D. Baker, Protein sequence optimization with a pairwise decomposable penalty for buried unsatisfied hydrogen bonds. Cold Spring Harbor Laboratory (2020), p. 2020.06.17.156646.

27. T. J. Brunette, F. Parmeggiani, P.-S. Huang, G. Bhabha, D. C. Ekiert, S. E. Tsutakawa, G. L. Hura, J. A. Tainer, D. Baker, Exploring the repeat protein universe through computational protein design. Nature. 528, 580–584 (2015).

28. J. R. Lydeard, B. A. Schulman, J. W. Harper, Building and remodelling Cullin-RING E3 ubiquitin ligases. EMBO Rep. 14, 1050–1061 (2013).

29. L. K. Langeberg, J. D. Scott, Signalling scaffolds and local organization of cellular behaviour. Nat. Rev. Mol. Cell Biol. 16, 232–244 (2015).

30. H. W. Schroeder Jr, L. Cavacini, Structure and function of immunoglobulins. J. Allergy Clin. Immunol. 125, S41–52 (2010).

31. P. Broz, V. M. Dixit, Inflammasomes: mechanism of assembly, regulation and signalling. Nat. Rev. Immunol. 16, 407–420 (2016).

32. L. Doyle, J. Hallinan, J. Bolduc, F. Parmeggiani, D. Baker, B. L. Stoddard, P. Bradley, Rational design of α-helical tandem repeat proteins with closed architectures. Nature. 528, 585–588 (2015).

33. I. Vulovic, Q. Yao, Y.-J. Park, A. Courbet, A. Norris, F. Busch, A. Sahasrabuddhe, H. Merten, D. D. Sahtoe, G. Ueda, J. A. Fallas, S. J. Weaver, Y. Hsia, R. A. Langan, A. Plückthun, V. H. Wysocki, D. Veesler, G. J. Jensen, D. Baker, Generation of ordered protein assemblies using rigid three-body fusion. Cold Spring Harbor Laboratory (2020), p. 2020.07.18.210294.

34. F. Khatib, S. Cooper, M. D. Tyka, K. Xu, I. Makedon, Z. Popovic, D. Baker, F. Players, Algorithm discovery by protein folding game players. Proc. Natl. Acad. Sci. U. S. A. 108, 18949–18953 (2011).

35. A. Chevalier, D.-A. Silva, G. J. Rocklin, D. R. Hicks, R. Vergara, P. Murapa, S. M. Bernard, L. Zhang, K.-H. Lam, G. Yao, C. D. Bahl, S.-I. Miyashita, I. Goreshnik, J. T. Fuller, M. T. Koday, C. M. Jenkins, T. Colvin, L. Carter, A. Bohn, C. M. Bryan, D. A. Fernández-Velasco, L. Stewart, M. Dong, X. Huang, R. Jin, I. A. Wilson, D. H. Fuller, D. Baker, Massively parallel de novo protein design for targeted therapeutics. Nature. 550, 74–79 (2017).

36. P. Hosseinzadeh, G. Bhardwaj, V. K. Mulligan, M. D. Shortridge, T. W. Craven, F. Pardo-Avila, S. A. Rettie, D. E. Kim, D.-A. Silva, Y. M. Ibrahim, I. K. Webb, J. R. Cort, J. N. Adkins, G. Varani, D. Baker, Comprehensive computational design of ordered peptide macrocycles. Science. 358, 1461–1466 (2017).

37. B. Dang, H. Wu, V. K. Mulligan, M. Mravic, Y. Wu, T. Lemmin, A. Ford, D.-A. Silva, D. Baker, W. F. DeGrado, De novo design of covalently constrained mesosize protein scaffolds with unique tertiary structures. Proc. Natl. Acad. Sci. U. S. A. 114, 10852–10857 (2017).

38. S. J. Fleishman, A. Leaver-Fay, J. E. Corn, E.-M. Strauch, S. D. Khare, N. Koga, J. Ashworth, P. Murphy, F. Richter, G. Lemmon, J. Meiler, D. Baker, RosettaScripts: a scripting language interface to the Rosetta macromolecular modeling suite. PLoS One. 6, e20161 (2011).

39. G. Bhardwaj, V. K. Mulligan, C. D. Bahl, J. M. Gilmore, P. J. Harvey, O. Cheneval, G. W. Buchko, S. V. S. R. K. Pulavarti, Q. Kaas, A. Eletsky, P.-S. Huang, W. A. Johnsen, P. J. Greisen, G. J. Rocklin, Y. Song, T. W. Linsky, A. Watkins, S. A. Rettie, X. Xu, L. P. Carter, R. Bonneau, J. M. Olson, E. Coutsias, C. E. Correnti, T. Szyperski, D. J. Craik, D. Baker, Accurate de novo design of hyperstable constrained peptides. Nature. 538, 329–335 (2016).

40. R. F. Alford, A. Leaver-Fay, J. R. Jeliazkov, M. J. O’Meara, F. P. DiMaio, H. Park, M. V. Shapovalov, P. D. Renfrew, V. K. Mulligan, K. Kappel, J. W. Labonte, M. S. Pacella, R. Bonneau, P. Bradley, R. L. Dunbrack Jr, R. Das, D. Baker, B. Kuhlman, T. Kortemme, J. J. Gray, The Rosetta All-Atom Energy Function for Macromolecular Modeling and Design. J. Chem. Theory Comput. 13, 3031–3048 (2017).

41. M. C. Lawrence, P. M. Colman, Shape complementarity at protein/protein interfaces. J. Mol. Biol. 234, 946–950 (1993).

42. B. Dang, M. Mravic, H. Hu, N. Schmidt, B. Mensa, W. F. DeGrado, SNAC-tag for sequence-specific chemical protein cleavage. Nat. Methods. 16, 319–322 (2019).

43. P. Virtanen, R. Gommers, T. E. Oliphant, M. Haberland, T. Reddy, D. Cournapeau, E. Burovski, P. Peterson, W. Weckesser, J. Bright, S. J. van der Walt, M. Brett, J. Wilson, K. J. Millman, N. Mayorov, A. R. J. Nelson, E. Jones, R. Kern, E. Larson, C. J. Carey, İ. Polat, Y. Feng, E. W. Moore, J. VanderPlas, D. Laxalde, J. Perktold, R. Cimrman, I. Henriksen, E. A. Quintero, C. R. Harris, A. M. Archibald, A. H. Ribeiro, F. Pedregosa, P. van Mulbregt, SciPy 1.0 Contributors, SciPy 1.0: fundamental algorithms for scientific computing in Python. Nat. Methods. 17, 261–272 (2020).

44. Z. L. VanAernum, F. Busch, B. J. Jones, M. Jia, Z. Chen, S. E. Boyken, A. Sahasrabuddhe, D. Baker, V. H. Wysocki, Rapid online buffer exchange for screening of proteins, protein complexes and cell lysates by native mass spectrometry. Nat. Protoc. 15, 1132–1157 (2020).

45. M. T. Marty, A. J. Baldwin, E. G. Marklund, G. K. A. Hochberg, J. L. P. Benesch, C. V. Robinson, Bayesian deconvolution of mass and ion mobility spectra: from binary interactions to polydisperse ensembles. Anal. Chem. 87, 4370–4376 (2015).

46. A. J. McCoy, R. W. Grosse-Kunstleve, P. D. Adams, M. D. Winn, L. C. Storoni, R. J. Read, Phaser crystallographic software. J. Appl. Crystallogr. 40, 658–674 (2007).

47. W. Kabsch, XDS. Acta Crystallogr. D Biol. Crystallogr. 66, 125–132 (2010).

48. Z. Otwinowski, W. Minor, in Methods in Enzymology (Academic Press, 1997), vol. 276, pp. 307–326.

49. M. D. Winn, C. C. Ballard, K. D. Cowtan, E. J. Dodson, P. Emsley, P. R. Evans, R. M. Keegan, E. B. Krissinel, A. G. W. Leslie, A. McCoy, S. J. McNicholas, G. N. Murshudov, N. S. Pannu, E. A. Potterton, H. R. Powell, R. J. Read, A. Vagin, K. S. Wilson, Overview of the CCP4 suite and current developments. Acta Crystallogr. D Biol. Crystallogr. 67, 235–242 (2011).

50. P. D. Adams, P. V. Afonine, G. Bunkóczi, V. B. Chen, I. W. Davis, N. Echols, J. J. Headd, L.-W. Hung, G. J. Kapral, R. W. Grosse-Kunstleve, Others, PHENIX: a comprehensive Python-based system for macromolecular structure solution. Acta Crystallogr. D Biol. Crystallogr. 66, 213–221 (2010).

51. G. N. Murshudov, A. A. Vagin, E. J. Dodson, Refinement of macromolecular structures by the maximum-likelihood method. Acta Crystallogr. D Biol. Crystallogr. 53, 240–255 (1997).

52. P. Emsley, K. Cowtan, Coot: model-building tools for molecular graphics. Acta Crystallogr. D Biol. Crystallogr. 60, 2126–2132 (2004).

53. B. L. Nannenga, M. G. Iadanza, B. S. Vollmar, T. Gonen, Overview of electron crystallography of membrane proteins: crystallization and screening strategies using negative stain electron microscopy. Curr. Protoc. Protein Sci. Chapter 17, Unit17.15 (2013).

54. T. Grant, A. Rohou, N. Grigorieff, cisTEM, user-friendly software for single-particle image processing. Elife. 7 (2018), doi:10.7554/eLife.35383.

55. A. Punjani, J. L. Rubinstein, D. J. Fleet, M. A. Brubaker, cryoSPARC: algorithms for rapid unsupervised cryo-EM structure determination. Nat. Methods. 14, 290–296 (2017).

