## supplementary_pdbs_scripts for "Reconfigurable asymmetric protein assemblies through implicit negative design": des3b_find_patches.py

```
import os
import pyrosetta
import argparse
import operator

pyrosetta.init('-mute all -ignore_unrecognized_res')

sfx_edge = pyrosetta.ScoreFunction()
sfx_edge.set_weight(pyrosetta.rosetta.core.scoring.ScoreType.hbond_lr_bb,1)

sfx = pyrosetta.get_fa_scorefxn()

parser=argparse.ArgumentParser()
parser.add_argument("pdbs", type=str, nargs="*")
parser.add_argument('-threshold', type=str, metavar='\b', default=- \
                    1.5, help='Only edges lower than threshold are considered; default -1.5')
parser.add_argument('-max_num_edges', type=str, metavar='\b', default=99999,
                    help='Not more than this number of edges are selected')
parser.add_argument('-apolar', action='store_true',
                        help='Only use hydrophobic pairs')
args=parser.parse_args()
max_num_edges = 1
if args.max_num_edges:
	max_num_edges = int(args.max_num_edges)

threshold = 0
if args.threshold:
	threshold = float(args.threshold)


def get_interface_edgestrand_energy(pose):
    '''Get interface edgestrand energy of binary complex (AB):
       pose'''
    separated_poses = []
    for chain in range(1,pose.num_chains()+1):
       separated_poses.append(pose.split_by_chain(chain))
    pose1 = separated_poses[0]
    pose2 = separated_poses[1]
    a = sfx_edge(pose1)
    b = sfx_edge(pose2)
    pose1.append_pose_by_jump(pose2,1)
    c = sfx_edge(pose1)
    edge_energy = ( c - (a + b))
    return edge_energy

def interface_residue_selector(pose):
    '''Fuction returning lists of interface residue chA and chB'''
    chain_A = pyrosetta.rosetta.core.select.residue_selector.ChainSelector("A")
    chain_B = pyrosetta.rosetta.core.select.residue_selector.ChainSelector("B")
    interfaceres_B = pyrosetta.rosetta.core.select.residue_selector.NeighborhoodResidueSelector(
        chain_A, 8, False)
    interfaceres_A = pyrosetta.rosetta.core.select.residue_selector.NeighborhoodResidueSelector(
        chain_B, 8, False)
    interfaceAlist = pyrosetta.rosetta.core.select.get_residues_from_subset(
        interfaceres_A.apply(pose))
    interfaceBlist = pyrosetta.rosetta.core.select.get_residues_from_subset(
        interfaceres_B.apply(pose))
    # print('\nInterface residues:')
    # print('InterfaceA: ',interfaceAlist)
    # print('InterfaceB: ',interfaceBlist)
    return interfaceAlist, interfaceBlist


def apolar_residues_interface(pose):
    '''Function returning lists of apolar interface residus
       chA and chB. Calls interface_residue_selector function'''
    hydrophobics = 'AILVMFYWP'
    interface = interface_residue_selector(pose)
    apolar_A = []
    apolar_B = []
    for i in interface[0]:
        if pose.residue(i).name1() in hydrophobics:
            apolar_A.append(i)
    for i in interface[1]:
        if pose.residue(i).name1() in hydrophobics:
            apolar_B.append(i)
    # print('\nApolar interface residues:')
    # print('Apolar interface A:', apolar_A)
    # print('Apolar interface B:', apolar_B)
    return apolar_A, apolar_B

def get_best_edges(pose, args):
    '''Returns dictionary of cross interface edges '''
    sfx = pyrosetta.get_fa_scorefxn()
    sfx(pose)
    if args.apolar:
        interfaceA = apolar_residues_interface(pose)[0]
        interfaceB = apolar_residues_interface(pose)[1]
    elif not args.apolar:
        interfaceA = interface_residue_selector(pose)[0]
        interfaceB = interface_residue_selector(pose)[1]
    edges = {}
    energies = pose.energies()
    graph = energies.energy_graph()
    for i in interfaceA:
        for j in interfaceB:
            try:
                interaction = graph.find_edge(i, j)
                interaction.fill_energy_map()
                val = interaction.dot(sfx.weights())
            except:
                pass
            else:
                if not val == 0.0:
                    edge = str(i)+'-'+str(j)
                    edges[edge] = interaction.dot(sfx.weights())
    best_edges = {}
    for i in sorted(list(edges.items()), key=operator.itemgetter(1))[0:max_num_edges]:
        if i[1] <= threshold:
            best_edges[str(i[0])] = str(i[1])
    return best_edges

scores = []
scores.append("stufff \n resA\tresB\tssA\tssB\tpatch_energy\tedgestrand_energy\tdescription")

for pdb in args.pdbs:
	pose = pyrosetta.io.pose_from_file(pdb)
	ss = pyrosetta.rosetta.core.scoring.dssp.Dssp(pose)
	edgestrand_energy = get_interface_edgestrand_energy(pose)
	sfx(pose)
	best_edges = get_best_edges(pose,args)
	for edge in list(best_edges):
		resA = int(edge.split("-")[0])
		resB = int(edge.split("-")[1])
		scores.append(f"""{resA}\t{resB}\t{ss.get_dssp_secstruct(resA)}\t{ss.get_dssp_secstruct(resB)}\t{best_edges[edge]}\t{edgestrand_energy}\t{pdb.split("/")[-1]}""")

with open("patchscore.sc","a+") as scorefile:
	for line in scores:
		scorefile.write(line + "\n")
```
