## supplementary_pdbs_scripts for "Reconfigurable asymmetric protein assemblies through implicit negative design": design_pipeline.txt

Interface design 

After generating complexes docked via their beta edgestrands(see Sahtoe et al 2021 for details on edge strand docking https://www.pnas.org/content/118/17/e2021569118), the interface can be designed using the following strategies.

Option 1: enforce polar interfaces using constraints

command to design the polar heterodimer interfaces:
    <path_to_rosettascripts> -s <input_dock> -holes:dalphaball <path_to_DAlphaBall.gcc> -corrections::beta_nov16 true -parser:protocol des1_polar_constraints.xml

DAlphaBall may be generated by navigating to Rosetta/main/source/external/DAlpahBall and typing make. Please make sure to put the compiler libraries in accessible directories, e.g. on Mac OSX, conda install -c anaconda gfortran_osx-64 gmp then ln -s ~/miniconda3/lib/libgfortran.* /usr/local/lib

Option 2: explicit hydrogen bond networks

To generate hydrogenbond networks across the interface:
    <path_to_rosettascripts> -s <input_dock> -holes:dalphaball <path_to_DAlphaBall.gcc> -beta -parser:protocol des2a_hbnet.xml 

Then design the rest of the interface while fixing the hydrogenbonds networks generated in des2a_hbnet.xml:
    <path_to_rosettascripts> -s <output_pdb_from_des2a> -holes:dalphaball <path_to_DAlphaBall.gcc> -corrections::beta_nov16 -parser:protocol des2b_design_around_hbnet.xml -parser:script_vars cst_file=<cst_file_genetated_by_des2a>


Option 3: polar interface design around hydrophobic patches

Initial interface design:
    <path_to_rosettascripts> -s <input_dock> -holes:dalphaball <path_to_DAlphaBall.gcc> -corrections::beta_nov16 true -parser:protocol des3a_np_design.xml

Then select the best hydrophobic interacting residue pairs across the interface:
    python des3b_find_patches.py <output_pdb_from_des3a> -apolar

The python script requires pyrosetta4 (https://www.rosettacommons.org/manuals/latest/all_else/d2/dfc/md_src_python__py_rosetta__r_e_a_d_m_e.html), python 3.6 and several standard python modules

From the resulting score file, select interaction pairs with patch_energy <= -3 & edgestrand_energy <= -5. For each selected interaction pair run script des3c_patch_design_layer.xml OR run script des3d_patch_design_comp.xml. 'resnums' are the residue numbers selected by des3b:
    <path_to_rosettascripts> -s <output_pdb_from_des3a> -holes:dalphaball <path_to_DAlphaBall.gcc> -corrections::beta_nov16 true -parser:protocol <des3c_or_des3d_xml> -parser:script_vars fix='resnums'


For options 1-3, filter final designs on: 

score_per_resA <= -2.5
score_per_resB <= -2.5
interface_sc >= 0.65   
packstat >= 0.65
vbuns <= 0
sbuns <= 1
rmsd_before_after_relax_rmsd <=  2
ddg_per_1000sasa <= -25
geometry == 1
ddg <= -40
ddg_hydrophobic <= -15


Generate rigid fusions:

The following example command rigidly fuses DHR53 to the N terminus of chain A of the input heterodimer:

<path_to_rosettascripts>
		-out::file::pdb_comments \
        -beta \
        -holes:dalphaball <path to dalphaball build> \
        -indexed_structure_store:fragment_store /net/databases/VALL_clustered/connect_chains/ss_grouped_vall_helix_shortLoop.h5 \
        -s input_dock.pdb \
        -parser:script_vars att_term='n_term' att_pdb='input_DHR53.pdb' dhr_rep_len='42' chain='A' loops='6' \

'dhr_rep_len' is the length of a repeat subunit for the input DHR, and 'loops' is the number of internal loops in chain A protomer of the heterodimer plus one. 

select fusions on

score_per_res <= -2.7
sc2 >= 0.65
sc2_int_area >= 500
buns_int <= 0
score_per_res_int <= -2.7
rmsd <= 1
geometry == 1
no_additional_contact <= 0
ss_count >= 5
ss_count2 >= 5
