## supplementary_pdbs_scripts for "Reconfigurable asymmetric protein assemblies through implicit negative design": readme.txt

All unfused monomers and single fusions (except 274A53 and 274B53) are models generated by the Rosetta design scripts described here, or variations thereof.

274A53 and 274B53 were generatred by grafting the interface of LHD274 onto the crystal structure of 29A53-29B53. Models of single fusions bound to unfused LHD274A or B were then generated by alignment in Pymol. 

Models of bivalent connectors, trivalent connectors, and oligomeric hubs were generated by the aignment along shared DHR repeats and subsequent splicing of sequences, either manually in Pymol or using a custom PyRosetta script. These models were generally not relaxed in Rosetta.

Models of assemblies were generated by alignment either manually in Pymol or using a custom PyRosetta script. These models were generally not relaxed in Rosetta.
